# Log Transformation Improves Dating of Phylogenies

**DOI:** 10.1101/2019.12.20.885582

**Authors:** Uyen Mai, Siavash Mirarab

## Abstract

Phylogenetic trees inferred from sequence data often have branch lengths measured in the expected number of substitutions and therefore, do not have divergence times estimated. These trees give an incomplete view of evolutionary histories since many applications of phylogenies require time trees. Many methods have been developed to convert the inferred branch lengths from substitution unit to time unit using calibration points, but none is universally accepted as they are challenged in both scalability and accuracy under complex models. Here, we introduce a new method that formulates dating as a non-convex optimization problem where the variance of log-transformed rate multipliers are minimized across the tree. On simulated and real data, we show that our method, wLogDate, is often more accurate than alternatives and is more robust to various model assumptions.

## 1 Introduction

Phylogenetic inference from sequence data does not reveal divergence time (i.e. exact timing of evolutionary events) unless paired with external timing information. Under standard models of sequence evolution, the evolutionary processes, including sequence divergence, are fully determined by the product of the absolute time and mutation rates in a non-identifiable form. Thus, these models measure branch lengths in the unit of expected numbers of mutations per site (since standard models like GTR [48] only allow substitutions, focusing on these models, we use *substitutions* and *mutations* interchangeably throughout this paper). Nevertheless, knowing divergence times is crucial for understanding evolutionary processes [10, 16] and is a fundamental need in many clinical applications of phylogenetics and phylodynamics [52]. A commonly used approach first infers a phylogeny with branch lengths in the unit of substitution per site and then dates the phylogeny by translating branch lengths from substitution unit to time unit; co-estimation of topology and dates is also possible [6] though its merits have been debated [**?**].

The fundamental challenge in dating is to find a way to factorize the number of substitutions into the product of the evolutionary rate and time. A common mechanism allowing this translation is to impose soft or hard constraints on the timing of *some* nodes of the tree, leaving the divergence times of the remaining nodes to be inferred based on the constrained nodes. Timing information is often in one of two forms: calibration points obtained from the geological record [23] and imposed on either internal nodes or tips that represent fossils [see 3], or tip sampling times for fast-evolving viruses and bacteria. The constraints still leave us with a need to extrapolate from observed times for a few nodes to the remaining nodes, a challenging task that requires a mathematical approach. Obtaining accurate timing information and formulating the right method of extrapolation are both challenging [e.g., see 35].

Many computational methods for dating phylogenies are available [e.g., see 24, 35], and a main point of differentiation between these methods is the clock model they assume [40]. Some methods rely on a strict molecular clock [56] where rates are effectively assumed to be constant [e.g., 26, 44, 50]. However, empirical evidence has now made it clear that rates can vary substantially, and ignoring these changes can lead to incorrect dating [**? ?**]. Consequently, there have been many attempts to *relax* the molecular clock and allow variations in rates. A main challenge in relaxing the clock is the need for a model of rates, and it is not clear what model should be preferred. As a result, many methods for dating using relaxed molecular clocks have been developed. Some of these methods allow rates to be drawn independently from a stationary distribution [1, 6, 51] while others model the evolution of rates with time [20] or allow correlated rates across branches [e.g., 5, 22, 28, 41, 45, 46, 49]. Despite these developments, strict molecular clocks continue to be used, especially in the context of intraspecific evolution where there is an expectation of relatively uniform rates [**?**].

Another distinction between methods is the use of explicit models [39]. Many dating methods use a parametric statistical model and formulate dating as estimating parameters in a maximum likelihood (ML) or Bayesian inference framework [e.g., 6, 26, 50, 51]. Another family of methods [e.g., 41, 46] formulate dating as optimization problems, including distance-based optimization [e.g., 54, 55], that avoid computing likelihood under an explicit statistical model. When the assumed parametric model is close to the reality, we expect parametric methods to perform well. However, these methods can be sensitive to model deviations, a problem that may be avoided by methods that avoid using specific models.

In this paper, we introduce LogDate, a new method of dating rooted phylogenies that allows variations in rates but without modeling rates using specific distributions. We define mutation rates necessary to compute time unit branch lengths as the product of a single global rate and a set of rate multipliers, one per branch. We seek to find the overall rate and all rate multipliers such that the log-transformed rate multipliers have the minimum variance. This formulation gives us a constrained optimization problem, which although not convex, can be solved in a scalable fashion using the standard approaches such as sequential least squares programming. While formulation of dating as an optimization problem is not new [26, 50], here we emphasize the realization that log-transforming the rate multipliers allows a wider variation in rates and results in more accurate dates. Our observation is in line with a recent change to RelTime [47] where the switch from arithmetic means to geometric means (between rates of sister lineages) has improved accuracy. In extensive simulation studies and three biological data sets, we show that a weighted version of LogDate, namely wLogDate, has higher accuracy in inferring node ages compared to alternative methods, including some that rely on time-consuming Bayesian inference. While wLogDate can date trees using both sampling times for leaves (e.g., in viral evolution) or estimated time of ancestors, most of our results are focused on cases with sampling times at the tips of the tree.

## 2 Methods

### 2.1 Definitions and notations

For a rooted binary tree *T* with *n* leaves, we give each node a unique index in [0,…, 2*n* − 2]. By convention, the root is always assigned 0, the other internal nodes are arbitrarily assigned indices in the range [1,…, *n* − 1], and the leaves are arbitrarily assigned indices in the range [*n* − 1,…, 2*n* − 2]. In the rest of this paper, we will refer to any node by its index. If a node *I* is not the root node, we let *par*(*i*) denote the parent of *i* and if *i* is not a leaf, we let *c*_*l*_(*i*) and *c*_*r*_(*i*) denote the left and right children of *i*, respectively. We refer to the edge connecting *par*(*i*) and *i* as *e*_*i*_.

We can measure each edge *e*_*i*_ of *T* in either time unit or substitution unit. Let *t*_*i*_ denote the divergence time of node *i*, i.e. the time when species *i* diverged into *c*_*l*_(*i*) and *c*_*r*_(*i*). Then for any node *i* other than the root, *τ*_*i*_ = *t*_*i*_ − *t*_*par*(*i*)_ is the length of the edge *e*_*i*_ in time unit. We measure divergence time of a node with respect to a fixed reference point in the past (i.e., time increases forward). Thus, we enforce *t*_*i*_ > *t*_*par*(*i*)_ for all *i*. Let *µ*_*i*_ be the substitution rate (per sequence site per time unit) on branch *e*_*i*_, then the expected number of substitutions per sequence site is *b*_*i*_ = *µ*_*i*_*τ*_*i*_. Let *τ* = [*τ*_1_,…, *τ*_2*n*−2_] and **b** = [*b*_1_,…, *b*_2*n*−2_].

From sequence data, **b** can be inferred using standard methods such as maximum parsimony [9], minimum evolution Rzhetsky and Nei [36], neighbor-joining [11, 37], and maximum-likelihood (ML) [8, 13, 31]. We let 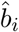 denote the estimate of *b*_*i*_ by an inference method and let 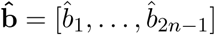.

In this paper, we are interested in computing *τ* from 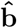. The computation of *τ* from 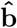 is complicated by two factors: (1) the possibility of change among rates, and (2) deviations of the inferred edge length 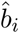 from the true value *b*_*i*_.

To better describe the mathematical formulation of the optimization problem, we first do the following change of variables. Assuming the mutation rates on the branches are distributed around a global rate *µ*, we define 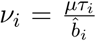. Let **x** = [*ν*_1_,…, *ν*_2*n*−2_, *µ*]; our goal of finding *τ* is identical to finding **x**.

### 2.2 Dating as a constrained optimization problem

We formulate dating as an optimization problem on 2*n* − 1 variables **x** = [*ν*_1_,…, *ν*_2*n*−2_, *µ*], subject to the linear constraints defined by calibration points and/or sampling times. Many existing methods, including LF [26] and LSD [50], can be described in this framework, with the choice of the objective function distinguishing them from each other. We start by describing the setup of the constraints enforced by a set of calibration points/sampling times, and show that they can all be written as linear equations on **x**. We then give the formulation of both LF and LSD in this framework and use their formulation to motivate our own new approach. Finally, we describe strategies to solve the wLogDate optimization problem.

#### 2.2.1 Linear constraints Ψ from sampling times

For any pair of nodes (*i, j*) (where each of *i* and *j* can either be a leaf or an internal node) with enforced divergence times (*t*_*i*_, *t*_*j*_), the following constraint *Ψ*(*i, j*) must be satisfied

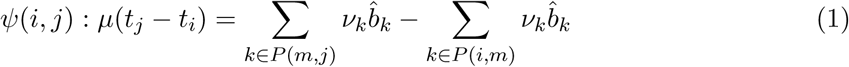

where *m* is the LCA of *i* and *j* and *P* (*m, j*) and *P* (*i, m*) are the paths connecting *m* to *j* and *I* to *m*, respectively. Thus, given *k* time points, *k*(*k* − 1)*/*2 constraints must hold. However, only *k* − 1 of these constraints imply all others, as we show below.

Let *t*_0_ be the *unknown* divergence time at the root of the tree. For *k* calibration points *t*_1_,…, *t*_*k*_, we can setup *k* constraints of the form:

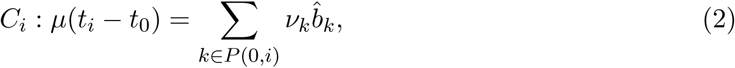

where node 0 is the root and *P* (0, *i*) is the path from the root to node *i*. For any pair (*i, j*), the linear constraint given in Eq. 1 can be derived by subtracting *C*_*i*_ from *C*_*j*_ side by side. Also, we can remove *t*_0_ from the set of constraints by subtracting *C*_1_ from all other constraints *C*_2_,…, *C*_*k*_. This gives us the final *k* − 1 linear constraints on **x**, which we denote as Ψ. We can build Ψ using Algorithm 1 (Supplementary material).

#### 2.2.2 Optimization Criteria

Since 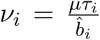, the distribution of *ν*_*i*_ is influenced by both the distribution of the rates (*µ*_*i*_) and the distribution of 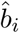 around *b*_*i*_. In traditional strict-clock models [56] such as those used by Least-Squares Dating (LSD) [50] and Langley-Fitch (LF) [26], a constant rate is assumed throughout the tree (∀_*i*_*µ*_*i*_ = *µ*). Under this model, the distribution of *ν*_*i*_ is determined by deviations of 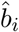 from *b*_*i*_.

Langley and Fitch [26] (LF) modeled the number of *observed* substitutions *per sequence site* on a branch *i* by a Poisson distribution with mean *λ* = *µτ*_*i*_ and modeled the total number of *observed* substitutions as 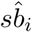 ∼ *Poisson*(*sµτ*_*i*_), where *s* is the sequence length. Therefore, by changing variable, we can write the log-likelihood function as:

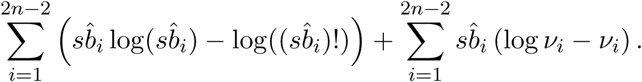

Given *s* and 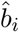, LF finds **x** that maximizes the log-likehiood function and subject to the constraints Ψ. As such,

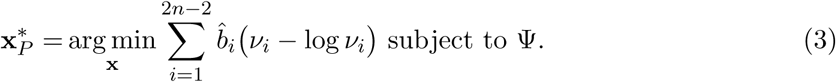

To *et al*. [50] assume 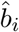 follows a Gaussian model: 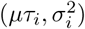 and assume the variance is approximated by 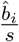 (the method includes smoothing strategies omitted here). Then, the negative log likelihood function can be written as:

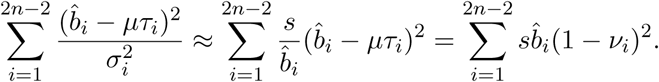

Thus, the ML estimate can be formulated as:

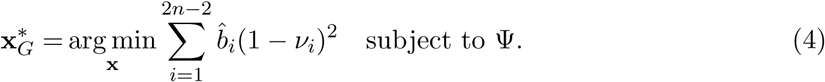

Both LF and LSD have convex formulations. Langley and Fitch [26] proved that their negative log-likelihood function is convex and thus the local minimum is also the global minimum. Our constraint-based formulation of LF also can be easily proved convex by showing its Hessian matrix is positive definite. To *et al*. [50] pointed out their objective function is a weighted least-squares. Using our formulation, we also see that Eq. 4 together with the calibration constraints form a standard convex quadratic optimization problem which has a unique analytical solution.

### 2.3 LogDate Method

LF and LSD only seek to model the errors in 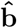and ignore true rate heterogeneity. Strict-clock assumption is now believed to be unrealistic in many settings [17, 32, 43], motivating relaxed clocks, typically by assuming that *µ*_*i*_s are drawn i.i.d. from some distribution [e.g., 1, 6, 51]. Most methods rely on presumed parametric distributions (typically, LogNormal, Exponential, or Gamma) and estimate parameters using ML [51], MAP [1], or MCMC [4, 6].

Our goal is to avoid explicit assumptions on the model that generates the rate multipliers. Instead, we follow the assumption shared by existing methods like LSD and LF: we assume that given two solutions of **x** both satisfying the calibration constraints, the solution with less variability in *ν*_*i*_ values is more preferable. Thus, we prefer solutions that minimize deviations from a strict clock while allowing deviations. However, to move away from strict clocks, we penalize rates in the log-space, and thus, allow for more variations than existing methods. A natural way to minimize deviations from the clock is to minimize the variance of 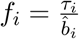. This can be achieved by finding *µ* and all *ν*_*i*_ such that *ν*_*i*_ is centered at 1 and 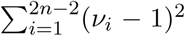 is minimized. Interestingly, the ML function used by LSD (Eq. 4) is a weighted version of this approach.

The minimum variance principle results in a fundamental asymmetry: multiplying or dividing the rate of a branch by the same factor is penalized differently (Fig 1a). For example, the penalty for *ν*_*i*_ = 4 is more than ten times larger than *ν*_*i*_ = 1*/*4. The LF model is more symmetrical than LSD but remains asymmetrical (Fig 1a). This asymmetry results from the asymmetric distribution of the Poisson distribution around its mean, especially for small mean, in log scale (Fig 1b). Because of this asymmetry, methods that only consider branch length estimation error judge a very small 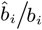to be within the realm of possible outcomes, and thus penalize *ν*_*i*_ < 1 multipliers less heavily than *ν*_*i*_ > 1.

**Figure 1:**
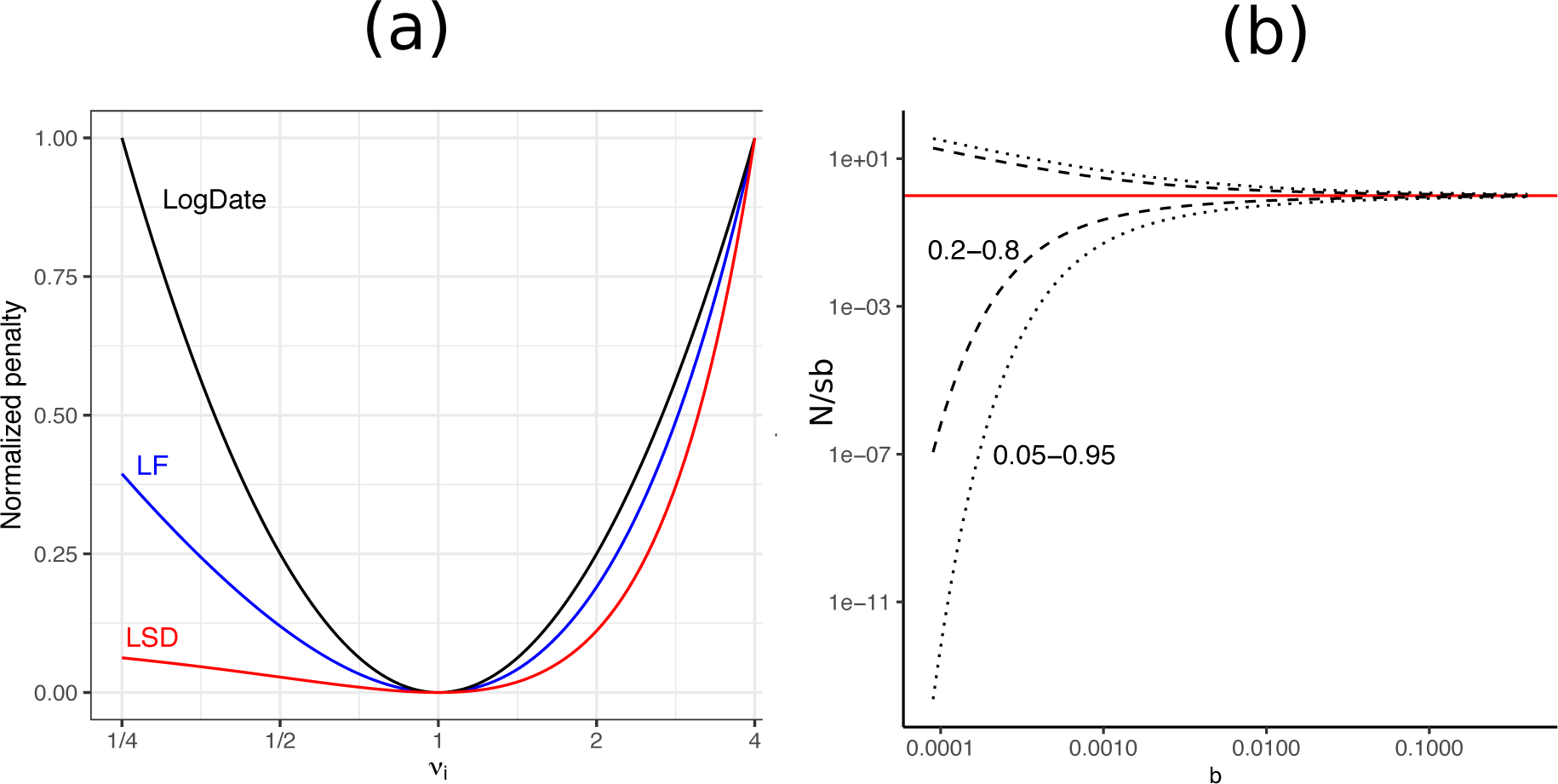
(a) The penalty associated to multiplying a single edge *i* with multiplier *ν*_*i*_ in LSD, LF, and LogDate approaches, as shown in Equations 3, 4, and 5. To allow comparison, we normalize the penalty to be zero at *ν* = 1 and to be 1 at *ν* = 4. (b) The confidence interval of the ratio between estimated and true branch length using the Poisson model. For this purpose of this exposition, we assume that the estimated branch length simply equals the number of substitutions occurring on the branch, which itself follows a Poisson distribution (as in the Jukes and Cantor [21] model), divided by sequence length (i.e., *N/s*). With this simplifying assumptions, the CI for 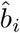 is between 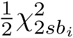 and 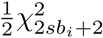; we draw the CI for 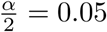and 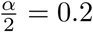 to get 0.2–0.8 and 0.05–0.095 intervals for 0.0001 ≤ *b*_*i*_ ≤ 0.4.

Our method is based on a fundamental assumption, which we call the *symmetry of ratios*: the penalty for multiplying a branch by a factor of *ν* should be no different than dividing the branch by *ν*. Note that this assertion is only applicable to true variations of the mutation rate (i.e, ignoring branch length estimation error). While this assumption is not motivated here by a specific probabilistic argument, we make the following case. If one considers the distribution of rate *multipliers* for various branches, absent of explicit model, it is reasonable to assume that the rates tend to increase by a factor of *ν* as often as they decrease by a factor of *ν* around the overall rate. Such an assumption would then motivate a method that penalizes *ν* and 1/*ν* identically. Objective functions of LSD and LF do not satisfy the symmetry of ratios. However, note that they do not seek to model true rate variation. Instead, they assume a strict clock and seek to model variations in estimated branch length. The usage of geometric means in the current version of RelTime [47], however, can be interpreted as a form of log-transformation. The success of the new version of RelTime over its predecessor further motivates the usage of log-transformation.

To ensure the symmetry of ratios, we propose taking the logarithm of the multipliers *ν*_*i*_ before minimizing their variance. Minimizing the variance of the rates in log-scale is the essence of our method. It achieves the symmetry, and, by changing the scale, it allows for much wider deviations from a strict clock that simply minimizing the variance.

#### 2.3.1 LogDate optimization function

We formulate the LogDate problem as follows. Given 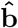 and the set of calibration constraints described earlier, we seek to find

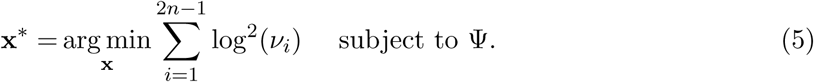

The objective function 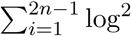(*ν*_*i*_) satisfies the symmetry of ratio property (Fig. 1a). Since *ν*_*i*_ values are multipliers of rates around *µ*, if we assume *µ* is the mean rate, the LogDate problem is equivalent to minimizing the variance of the log-transformed rate multipliers (around their mean 1). The objective function only depends on *ν*_*i*_; however, note that *µ* is still included in the constraints and therefore is part of the optimization problem. This setting reduces the complexity of the objective function and speeds up the numerical search for the optimal solution. Since the values of *ν*_*i*_ close to 1 are preferred in Eq. 5, the optimal solution would push *µ* to the mean rate.

We can prove that if *ν*_*i*_ is drawn from a LogNormal distribution with *any parameters* that result in mode 1 (which is a reasonable condition for rate *multipliers*), the LogDate optimization problem is equivalent to finding *ν*_*i*_ that maximize the joint probability under the LogNormal model, subject to the constraints Ψ. A formal proof is given in Claim 1 (Supplementary materials).

The simple LogDate formulation, however, has a limitation: by allowing rates to vary freely in a multiplicative way, it fails to deal with the varied levels of relative branch error; i.e., the ratio of the estimated branch length to the true branch length 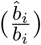. As 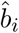 is estimated from the sequences, the error of 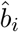 is directly related to the variations in the number of substitutions occurred along the branch *b*_*i*_. Let us assume sequences follow the Jukes and Cantor [21] model, and le *N*_*i*_ be the total number of substitutions occurred along branch *i* on a sequence with length *s*. Under Juke-Cantor model, we have *N*_*i*_ ∼ *Poisson*(*sµτ*_*i*_) and therefore, *var*(*N*_*i*_) = *sµτ*_*i*_. Therefore, the variance of the *expected number of substitutions* around the true branch length is 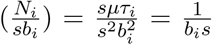. As Figure 1b shows, when *b*_*i*_ is small, 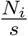 can easily vary by several orders of magnitude around *b*_*i*_. Furthermore, the distribution is not symmetric: drawing values several factors smaller than the mean is more likely than drawing values above the mean by the same factor. These analyses predict that the distribution of 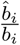 depends strongly on *b*_*i*_ – with smaller *b*_*i*_ giving higher variance - and is not symmetric.

The variances of the relative error 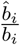 is difficult to compute analytically due to the involvement of the sequence substitution model and the method to estimate 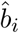, which are both unknown. Therefore, we instead use empirical analyses of the estimated branch lengths by PhyML to demonstrate our arguments. Consistent with our prediction, Figures S5 a and c illustrate that the relative error 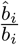 varies more in small branches and the distribution is not symmetric. These properties of the branch length estimates are not modeled in our LogDate formulation and we seek to incorporate them in a refined version of LogDate which will be described below.

Since the true branch length *b*_*i*_ is unknown, a common practice is to use the estimated 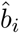 in place of *b*_i_ to estimate its variance as 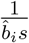. This explains why both LF and LSD objective functions (Eqs. 3 and 4) have a weight of 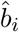 for each term of *ν*_*i*_. Following the same strategy, we propose weighting each log^2^(*ν*_*i*_) term in a way that reduces the contribution of short branches to the total penalty, and thus allows more deviations in the log space if the branch is small (and is thus subject to higher error). Since we log-transform *ν*_*i*_ and pursue a model-free approach, explicitly computing the weights to cancel out the variations of relative error among the branches is challenging. However, since the weights should reflect the variance of 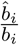 (logarithmic scale), they should monotonically increase with 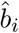 (Fig. 1b) to allow more variance for the *relative* errors in short branches than in long branches. We use 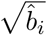 as weights, a selection driven by simplicity and empirical performance (shown in a later section).

The shortest branches require even more care. When the branch is very short, for a limited-size alignment, the evolution produces zero mutations with high probability. For these no-event branches, tree estimation tools report arbitrary small lengths (see Fig. S5), rendering 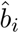 values meaningless for very small branches. To deal with this challenge, the r8s’s implementation of LF [42] collapses all branches with length 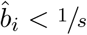<1/*s*. To *et al*. [50] proposed adding a smoothing constant *c/s* to each 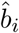 to estimate the variance of 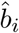, where *c* is a parameter that the user can tune. Following a similar strategy, we propose adding a small constant 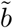 to each 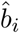 We choose 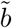 to be the maximum branch length that produces no substitutions with probability at least 1 − *α* for *α* ∈ [0, 1]. Recall that *N* is the total number of *actual* substitutions on a branch. Under the Jukes and Cantor [21] model, it is easy to show that arg 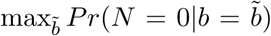 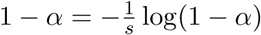. We choose this value as 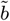 and set *α* = 0.01 by default. Thus, we define the weighted LogDate (wLogDate) as follows:

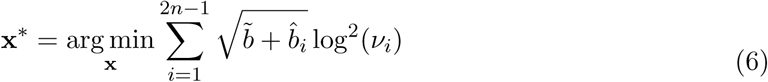

subject to Ψ.

#### 2.3.2 Solving the optimization problem

Both LogDate and wLogDate problems (Eq. 5 and Eq. 6) are non-convex, and hence solving them is non-trivial. The problem is convex if 0 ≤ *ν*_*i*_ ≤ *e*. For small clock deviation and small estimation error in *b*_*i*_, the *ν*_*i*_ values should be small so that the problem becomes convex with one local minimum. However, as *ν*_*i*_ ≤ *e* is not guaranteed, we have to rely on gradient-based numerical methods to search for multiple local minima and select the best solution we can find. To search for local minima, we use the Scipy solver with `trust-constr` [25] method. To help the solver work efficiently, we incorporate three techniques that we next describe.

**Computing Jacobian and Hessian matrices** analytically helps speedup the search. By taking the partial derivative of each *ν*_*i*_, we can compute the Jacobian, *J*, of Eq. 6. Also, since Eq. 6 is separable, its Hessian *H* is a (2*n* − 2) × (2*n* − 2) diagonal matrix. Simple derivations give us:

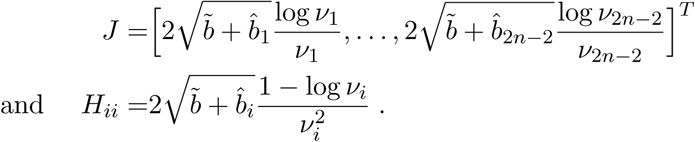

**Sparse matrix representation**further saves space and computational time. The Hessian matrix is diagonal, allowing us to store only the diagonal elements. In addition, the constraint matrix defined by Ψ is highly sparse. If all sampling times are given at the leaves, the number of non-zero elements in our (*n* − 1) × (2*n* − 1) matrix is *O*(*n* log *n*) (Claim 3; Supplementary materials). If the tree is either caterpillar or balanced, the number of non-zeroes reduced to Θ(*n*). Thus, we use sparse matrix representation implemented in the Scipy package. This significantly reduces the running time of LogDate.

**Starting from multiple feasible initial points** is necessary given that our optimization problem is non-convex. Providing initial points that are feasible (i.e. satisfied the calibration constraints) helps the SciPy solver work efficiently. We designed a heuristic strategy to find multiple initial points given sampling times *t*_1_,…, *t*_*n*_ of all the leaves (as is common in phylodynamics).

We first describe the process to get a single initial point. We compute the root age *t*_0_ and *µ* using root-to-tip regression (RTT) [44]. Next, we scale all branches of *T* to conform with Ψ as follow: let *m* = arg min_*i*_ *t*_*i*_ (breaking ties arbitrarily). Let *d*(*r, i*) denote the distance from the root *r* to node *i* and *P* (*r, m*) denote the path from *r* to *m*. For each node *i* in *P* (*r, m*), we set 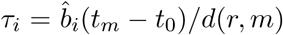. Then going upward from *m* to *r* following *P* (*m, r*), for each edge (*i, j*) we compute *t*_*j*_ = *t*_*i*_ − *τ*_*i*_ and recursively apply the process on the clade *i*. At the root, we set *t*_*m*_ to the second oldest (second minimum) sampling time and repeat the process on a new path until all leaves are processed. Since RTT gives us *µ*, to find *ν* we simply set 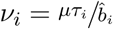.

To find multiple initial points, we repeatedly apply RTT to a set of randomly selected clades of *T* and scale each clade using the aforementioned strategy. Specifically, we randomly select a set *S* of some internal nodes in the tree and add the root to *S*. Then, by a post-order traversal, we visit each *u* ∈ *S* and date the clade *u* using the scaling strategy described above. We then remove the entire clade *u* from the tree but keep the node *u* as a leaf (note that the age of *u* is already computed) and repeat the process for the next node in *S*. The root will be the last node to be visited. After visiting the root, we have all the *τ*_*i*_ for all *i*. After having all the branches in time unit, we find **x** to minimize either Eq. 5 or Eq. 6, depending on whether LogDate or wLogDate is chosen. In a tree of *n* leaves, we have 2^(*n*−1)^ − 1 ways to select the initial non-empty set *S*, giving us enough room for randomization.

#### 2.3.3 Computing confidence interval

With the ability of wLogDate to work on any combination of sampling times/calibration points on both leaves and internal nodes (as long as at least two time points are provided), we design a method to estimate the confidence intervals for the estimates of wLogDate. We subsample the sampling times/calibration points given to us repeatedly to create *N* replicate datasets (where *N* is 100 by default, but can be adjusted). Note our subsampling is not exactly a bootstrapping procedure as node sampling times cannot be resampled with replacement. We then compute the time tree for each replicate to obtain *N* different estimates for the divergence time of each node, from which we can compute their confidence intervals (95% as default). This sampling would work best when we have a fairly large number of calibration points, which is the case in phylodynamic settings where all (or nearly all) sampling times for the leaves are given, or in large phylogenies where abundant calibration points can be obtained from fossils.

### 2.4 Experiments on simulated data

#### 2.4.1 Phylodynamics setting

To *et al*. [50] simulated a dataset of HIV *env* gene. Their time trees were generated based on a birth-death model with periodic sampling times. There are four tree models, namely D995 11 10 (M1), D995 3 25 (M2), D750 11 10 (M3), and D750 3 25 (M4), each of which has 100 replicates for a total of 400 different tree topologies. M1 and M2 simulate intra-host HIV evolution and are ladder-like while M3 and M4 simulate inter-host evolution and are balanced. Also, M4 has much higher root-to-tip distance (mean: 57) compared to M1–M3 (22, 33, and 29). Starting from conditions simulated by To *et al*. [50], we use the provided time tree to simulate the clock deviations. Using an uncorrelated model of the rates, we draw each rate from one of three different distributions, each of which is centered at the value *µ* = 0.006 as in To *et al*. [50]. Thus, we set each *µ*_*i*_ to *x*_*i*_*µ* where *x*_*i*_ is drawn from one of three distributions: LogNormal (mean:1.0, std: 0.4), Gamma (*α* = *β* = 6.05), and Exponential (*λ* = 1). Sequences of length 1000 were simulated for each of the model conditions using SeqGen [33] under the same settings as To *et al*. [50].

#### 2.4.2 Calibrations on autocorrelated rate model

We used the software NELSI and the same protocol as in [19] to simulate a dataset where the rates are autocorrelated. The dataset has 10 replicates, each contains 50 taxa. The time trees were generated under Birth-death model and the rate heterogeneity through time is modeled by the autocorrelation model ([22]) with the initial rate set to 0.01 and the autocorrelated parameter set to 0.3. DNA sequences (1000 bases) were generated under Jukes-Cantor model. We used PhyML Guindon *et al*. [13] to estimate the branch lengths in substitution unit from the simulated sequences while keeping the true topology. These trees are the inputs to wLogDate, RelTime, and LF to infer time trees.

### 2.5 Real biological data

#### H1N1 2009 pandemic

We re-analyze the H1N1 biological data provided by To *et al*. [50] which includes 892 H1N1pdm09 sequences collected worldwide between 13 March 2009 and 9 June 2011. We reuse the estimated PhyML [13] trees, 100 bootstrap replicates, and all the results of the dating methods other than LogDate that are provided by To *et al*. [50].

#### San Diego HIV

We study a dataset of 926 HIV-1 subtype B *pol* sequences obtained in San Diego between 1996 and 2018 as part of the PIRC study. We use IQTree [31] to infer a tree under the GTR+Γ model, root the tree on 22 outgroups, then remove the outgroups. Because of the size, we could not run BEAST.

#### West African Ebola epidemic

We study the dataset of Zaire Ebola virus from Africa, which includes 1,610 near-full length genomes sampled between 17 March 2014 and 24 October 2015. The data was collected and analyzed by Dudas *et al*. [7] using BEAST and re-analyzed by Volz and Frost [51] using IQTree to estimate the ML tree and *treedater* to infer node ages. We run LSD, LF, and wLogDate on the IQTree from Volz and Frost [51] and use the BEAST trees from Dudas *et al*. [7], which include 1000 sampled trees (BEAST-1000) and the Maximum clade credibility tree (BEAST-MCC). To root the IQTree, we search for the rooting position that minimizes the triplet distance [38] between the IQTree and the BEAST-MCC tree.

#### 2.5.1 Methods Compared

For the phylodynamics data, we compared wLogDate to three other methods: LSD [50], LF [26], and BEAST [4]. For all methods, we fixed the true rooted tree topology and only inferred branch lengths. For LSD, LF, and wLogDate, we used phyML [13] to estimate the branch lengths in substitution unit from sequence alignments and used each of them to infer the time tree. LSD was run in the same settings as the QPD* mode described in the original paper [50]. LF was run using the implementation in r8s [42]. wLogDate was run with 10 feasible starting points. For the Bayesian method BEAST, we also fixed the true rooted tree topology and only inferred node ages. Following To *et al*. [50], we ran BEAST using HKY+Γ8 and coalescent with constant population size tree prior. We used two clock models on the rate parameter: the strict-clock (i.e. fixed rate) model and the LogNormal model. For the strict-clock prior, we set clock rate prior to a uniform distribution between 0 and 1. For the LogNormal prior, we set the ucld.mean prior to a uniform distribution between 0 and 1, and ucld.stdev prior to an exponential distribution with parameter 1*/*3 (default). We always set the length of the MCMC chain to 10^7^ generations, burn-in to 10%, and sampling to every 10^4^ generations (identical to To *et al*. [50]).

For the autocorrelated rate model, we compared wLogDate to LF and RelTime [47], which is one of the state-of-the-art model-free dating methods. We randomly chose subsets of the internal nodes (10% on average) as calibration points and created 20 tests for each of the 10 replicates (for a total of 200 tests).

#### 2.5.2 Evaluation Criteria

On the simulated phylodynamics dataset where the ground truth is known, we compare the *accuracy* of the methods using several metrics. We compute the root-mean-square error (RMSE) of the true and estimated vector of the divergence times (*τ*) and normalize it by tree height. We also rank methods by RMSE rounded to two decimal digits (to avoid different ranks when errors are similar). In addition, we examine the inferred divergence time of the Most Recent Common Ancestor (tMRCA) and mutation rate. The comparison of methods mostly focuses on point-estimates of these parameters and the accuracy of the estimates (as opposed to their variance). In one analysis, we also compare the confidence intervals produced by wLogDate and BEAST on one model condition (M3 with LogNormal rate distribution). Finally, we examine the correlation between variance of the error in wLogDate and divergence times and branch lengths.

On the simulated data with autocorrelated rate, we show the distributions of the divergence times estimated by wLogDate, LF, and RelTime and report the RMSE normalized by tree height for each replicate.

On real data, we show lineage-through-time (LTT) plots [30], which trace the number of lineages at any point in time and compare tMRCA times to the values reported in the literature. We also compare the runtime of wLogDate to all other methods in all analyses.

## 3. Results

### 3.1 Simulated data for phylodynamics

We first evaluate the convergence of the ScipPy solver across 10 starting points (Fig. S1a). LogDate and wLogDate converge to a stable result after 50–200 iterations, depending on the model condition. Convergence seems easier when rates are Gamma or LogDate and harder when the rates are Exponential. Next, to control for the effect of the starting points on the accuracy of our method, we compare the error of these starting points with the wLogDate optimal point (Fig. S1b). In all model conditions, the optimal point shows dramatic improvement in accuracy compared to the starting point. We then compare different weighting strategies for LogDate (Table S4). In all model conditions, the weighting 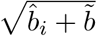 is one of the two best, so it is chosen as the default weighting for wLogDate. Moreover, wLogDate is never worse than LogDate, and under exponential clock models, appropriate weighting results in dramatic improvements (Table S4).

Next, we study the properties of wLogDate estimates in relation to: (1) the age of the node (Fig. 2a), (2) the length of the true branch in time unit (Fig. 2b), and (3) the error of the branch lengths (in substitution unit) estimated by PhyML (Fig. S4). Overall, we do not observe a substantial change in the mean estimation error of wLogDate as the node age and the branch length change. The variance, however, can vary with node ages (Figure 2a), especially in M3 and M4 model conditions. Moreover, longer branches have a tendency to have higher variance in absolute terms (Fig. 2b). However, note that the relative error (i.e., log-odds error) dramatically *reduces* as branches become longer (Fig. S4). In studying the effect of the error in branch length estimation, we see that wLogDate can underestimate the branch time if the branch length in substitution unit is extremely underestimated (Fig.S4a, Supplementary Materials). In some cases wLogDate under-estimates branch times by two order of magnitude or more; all of these cases correspond to super-short branches with substitution unit branch length under-estimated by three or four orders of magnitude (Fig.S4a). As mentioned previously, extremely short estimated branch lengths are often the zero-event branches (Fig.S5), which are unavoidable for short sequences.

**Figure 2:**
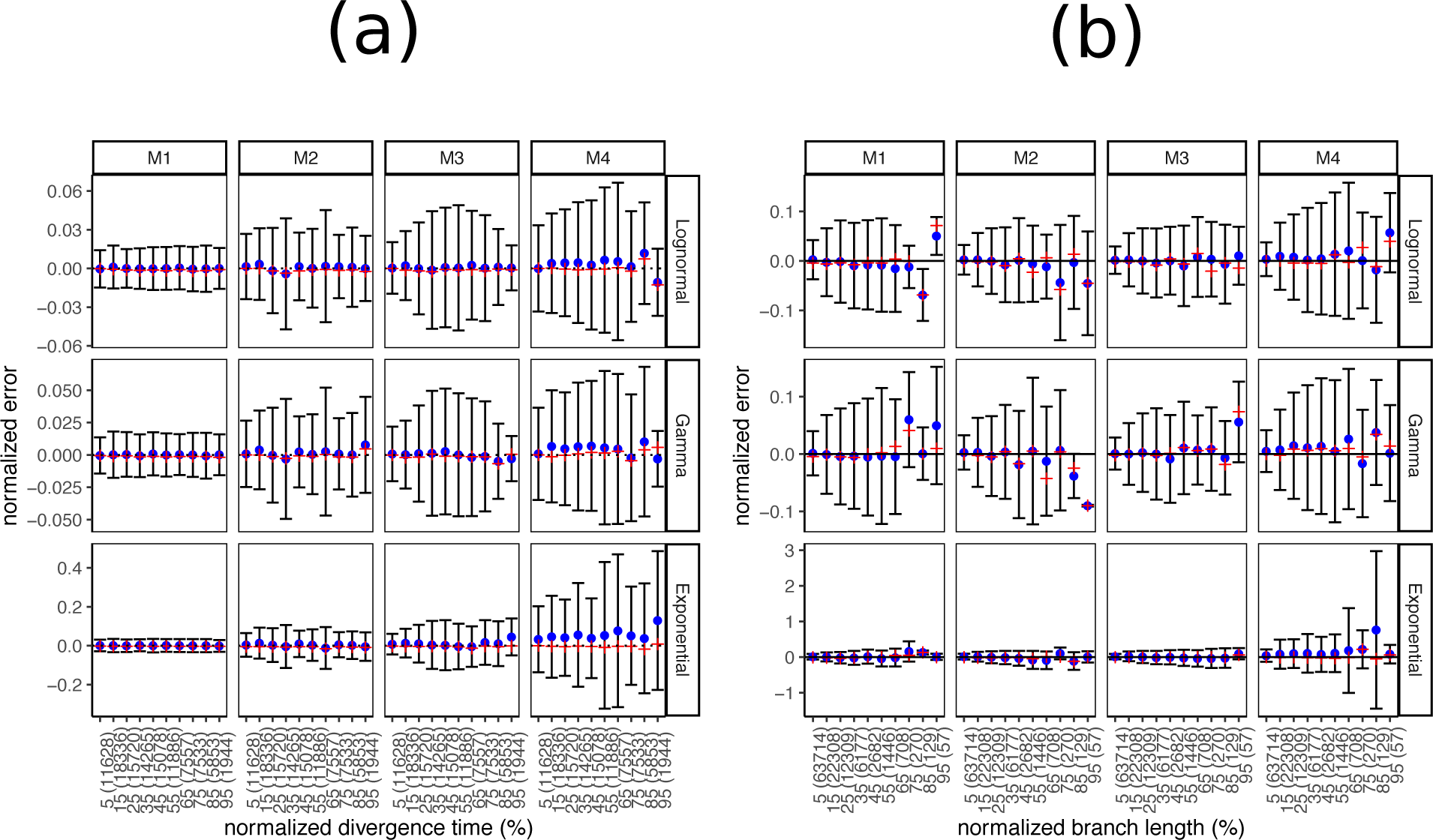
Analyses of wLogDate on inferring branch lengths on simulated data. (a) error normalized by tree height versus divergence time (i.e. the time of the midpoint of each branch); both axes are normalized by the tree height. (b) error versus branch length (in time unit); both axes are normalized by the maximum branch length. For both (a) and (b), the x-axis is discretized into 10 bins of equal size. We label the bins by their median values, relative to either the tree height for (a) or the maximum branch length for (b). We also show the number of points in each bin in parentheses. Note the small number of points in the final bins in panel (b). For each bin, the blue dot represents the mean, the red cross represents the median, and the bar represents one standard deviations around the mean.

We next compare wLogDate to alternative methods, namely LF, LSD, and BEAST with strict-clock and Lognormal clock. Measured by RMSE, the accuracy of all methods varies substantially across model trees (M1 – M4) and models of rate variation (Fig. 3). Comparing methods, for many conditions, wLogDate has the lowest error, and in many others, it is ranked second best (Table 1). Across all conditions, wLogDate has a mean rank of 1.75, followed by BEAST with strict clock with a mean rank 2; mean normalized RMSE of wLogDate, LF, BEAST-strict, BEAST-LogNormal, and LSD are 0.072, 0.074, 0.077, 0.087, and 0.116, respectively. Interestingly, in contrast to wLogDate, LSD seems to often underestimate branch times for many short branches even when they are estimated relatively accurately in substitution units (Fig. S4b, Supplementary Materials). For all methods, errors are an order of magnitude smaller for the LogNormal and Gamma models of rate variations compared to the Exponential model. In terms of trees, M4, which simulates inter-host evolution and high the largest height, presents the most challenging case for all methods. Interestingly, wLogDate has the best accuracy under all parameters of M4 tree and also all parameters of M3 (thus, both inter-host conditions). On M1, all methods have very low error and perform similarly (Fig. 3).

**Table 1:**
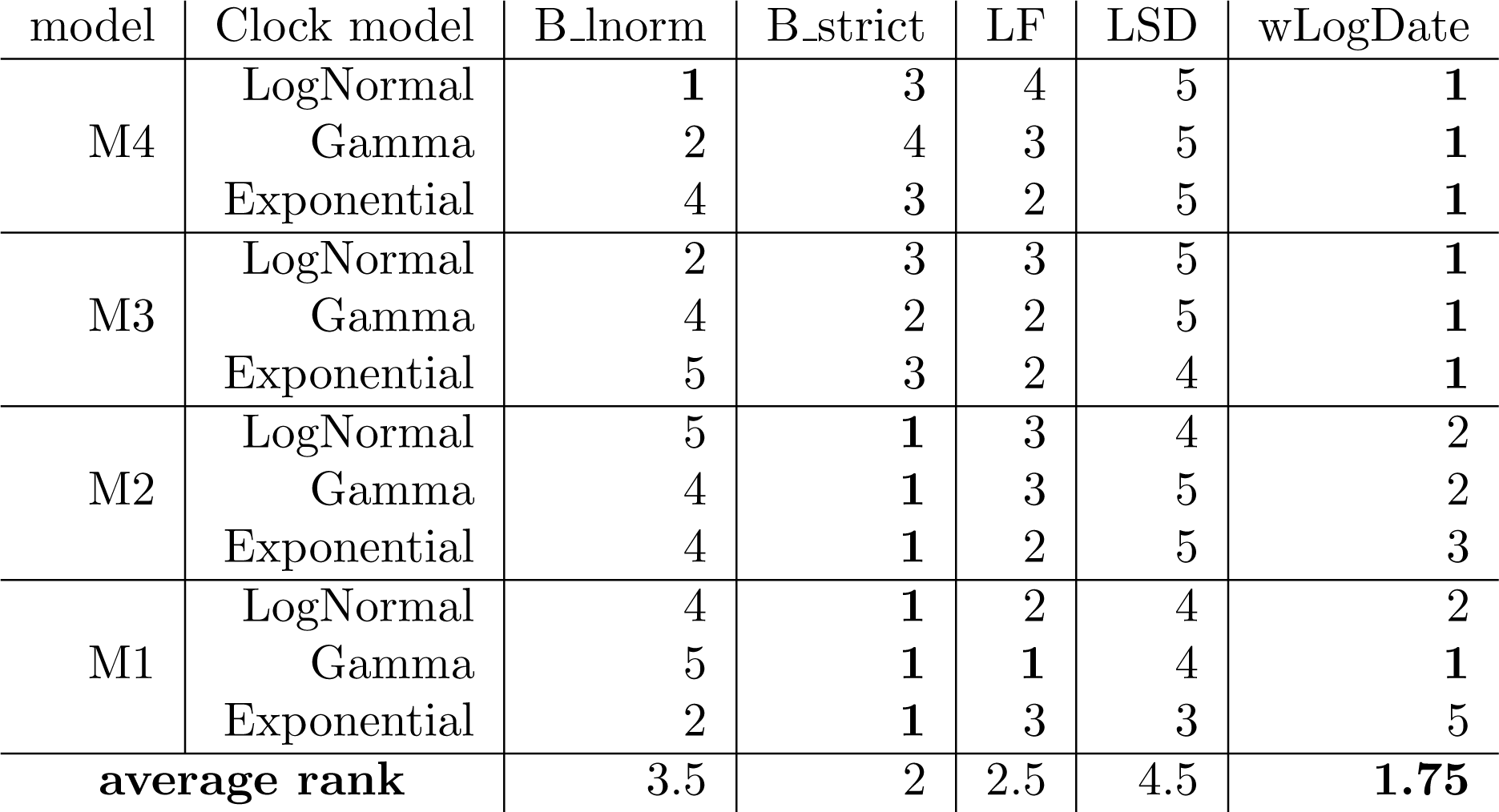
Ranking of the dating methods under different model conditions. For each model condition, the average RMSE of all internal node ages is computed and ranked among the dating methods (rounded to two decimal digits). The best method is shown in bold.

**Figure 3:**
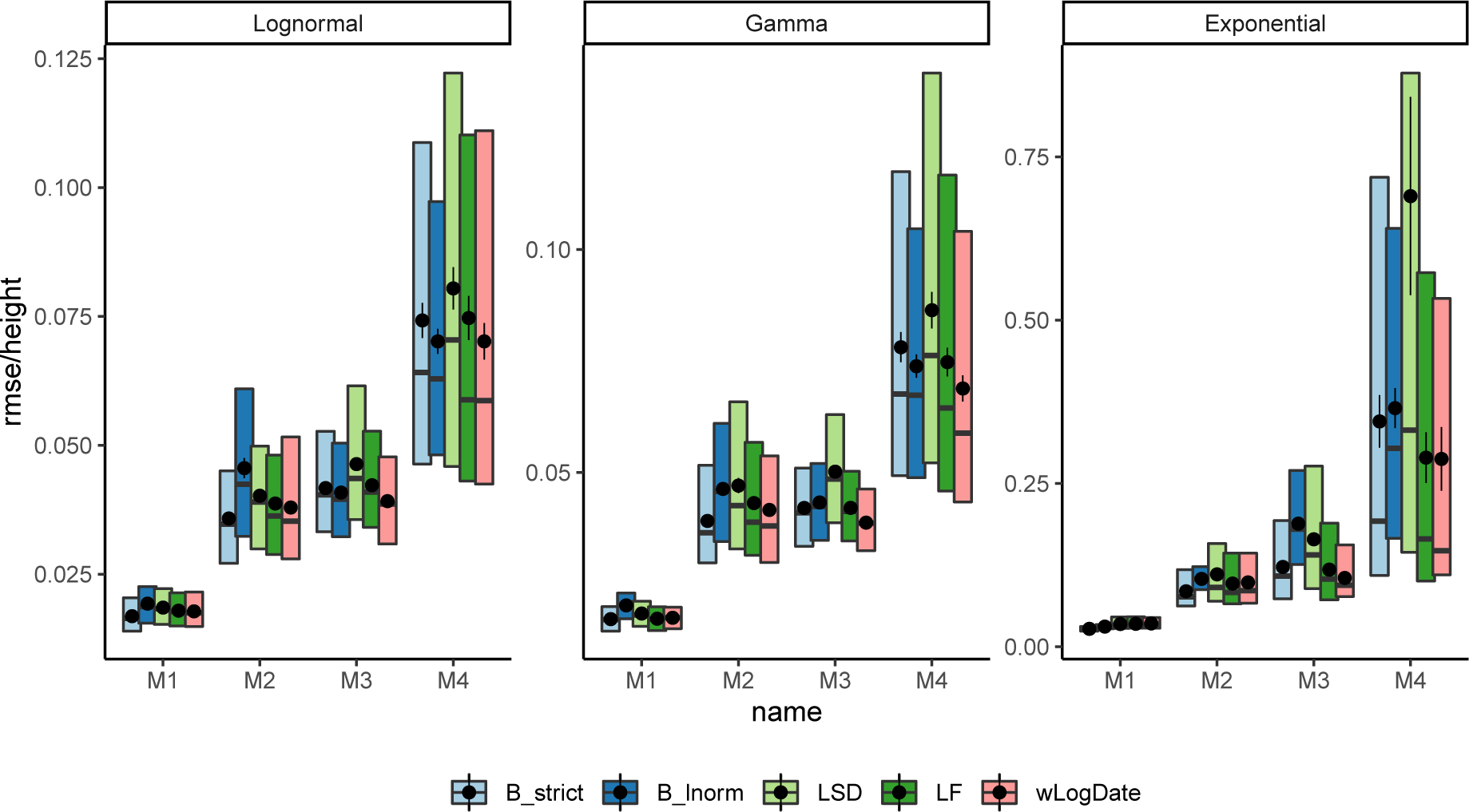
Distributions of RMSE normalized by the tree height for internal node ages inferred by all methods on model trees M1–M4, each with clock models Lognorm, Gamma, and Exponential. Boxes show median, 10% and 90% quantiles; dots and error bars show mean and standard error (100 replicates).

Among other methods, results are consistent with the literature. Despite its conceptual similarity to wLogDate, LSD has the worst accuracy. On M1 and M2, LSD is competitive with other methods; however, on M3 and M4, it has a much higher error, especially with the Exponential model of rate variation. With the LogNormal clock model, BEAST-LogNormal is better than BEAST-strict only for M4 but not for M1–M3; in fact, BEAST-LogNormal has the highest error for the M2 condition. This result is surprising given the correct model specification. Nevertheless, BEAST-LogNormal is competitive only under the LogNormal model of rate variation and is one of the two worst methods elsewhere. Thus, BEAST-LogNormal is sensitive to model misspecification. In contrast, BEAST-strict is less sensitive to the model of rate variation and ranks among the top three in most cases. In particular, BEAST-strict is always the best method for intra-host ladder-like trees M1 and M2.

Accuracy of tMRCA follows similar patterns (Fig. 4). Again, the Exponential rate variation model is the most difficult case for all methods, resulting in biased results and highly variable error rates for most methods. In all conditions of M3 and M4, wLogDate has the best accuracy and improves on the second best method by 9 – 66% (Table 2). For M1 and M2, BEAST-strict is often the best method. The mean tMRCA error of wLogDate across all conditions is 4.83 (years), which is substantially better than the second best method, BEAST-strict (6.21).

**Table 2:** Mean absolute error of the inferred tMRCA of BEAST strict, BEAST lognorm, LF, LSD, RTT, and wLogDate. For wLogDate, parenthetically, we compare it with the best (↑) or second best (↓) method for each condition. We show percent improvement by wLogDate, as measured by the increase in the error of the second best method (wLogDate or the alternative) divided by the error of the best method.

| Tree | Clock Model | B_strict | B_lnorm | LF | LSD | RTT | wLogDate |
| --- | --- | --- | --- | --- | --- | --- | --- |
| M4 | Lognormal | 6.99 | 9.50 | 6.66 | 7.38 | 9.28 | **6.11** ( 9% ↓) |
|  | Gamma | 7.83 | 10.48 | 7.02 | 8.48 | 8.24 | **6.28** (12% ↓) |
|  | Exponential | 43.5 | 140.9 | 116.2 | 62.2 | **31.5** | 32.5 (3% ↑) |
| M3 | Lognormal | 1.37 | 2.60 | 1.21 | 1.39 | 1.46 | **1.03** (17% ↓) |
|  | Gamma | 1.60 | 3.14 | 1.23 | 1.67 | 1.42 | **0.97** (27% ↓) |
|  | Exponential | 5.76 | 34.67 | 4.87 | 8.35 | 3.39 | **2.94** (66% ↓) |
| M2 | Lognormal | **1.40** | 1.41 | 1.50 | 1.63 | 2.19 | 1.47 ( 5% ↑) |
|  | Gamma | 1.54 | **1.44** | 1.75 | 1.92 | 2.56 | 1.66 (15% ↑) |
|  | Exponential | **3.39** | 4.59 | 4.28 | 5.27 | 5.23 | 3.72 ( 10% ↑) |
| M1 | Lognormal | **0.28** | **0.28** | 0.30 | 0.37 | 0.78 | 0.30 ( 7% ↑) |
|  | Gamma | **0.27** | 0.29 | 0.32 | 0.35 | 0.80 | 0.30 (11% ↑) |
|  | Exponential | **0.60** | 1.11 | 0.79 | 0.82 | 1.37 | 0.69 (15% ↑) |
| **Average** |  | 6.21 | 17.54 | 12.17 | 8.13 | 5.68 | **4.83** |

**Figure 4:**
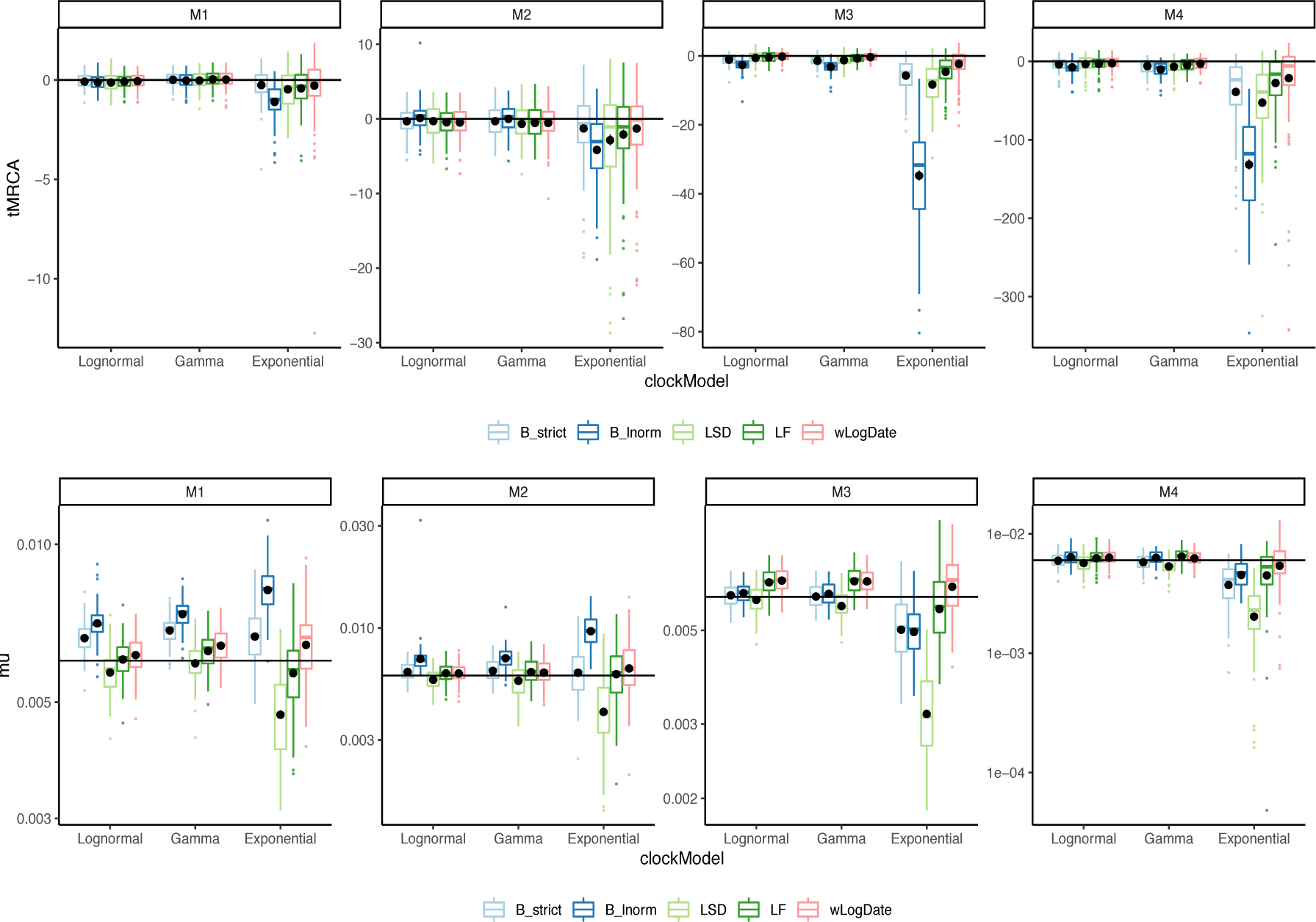
The inferred (top) tMRCA and (bottom) expected mutation rate on different tree models and clock models. Distributions are over 100 replicates. The solid horizontal lines indicate the true mutation rate and tMRCA. Each black is the average of the inferred values for each method under each model condition. We remove 6 outlier data points (2 LF, 1 LSD, 2 BEAST-LogNormal, 1 BEAST-Strict) with exceptional incorrect tMRCA (< −350) in the M4/Exponential model.

**Figure 5:**
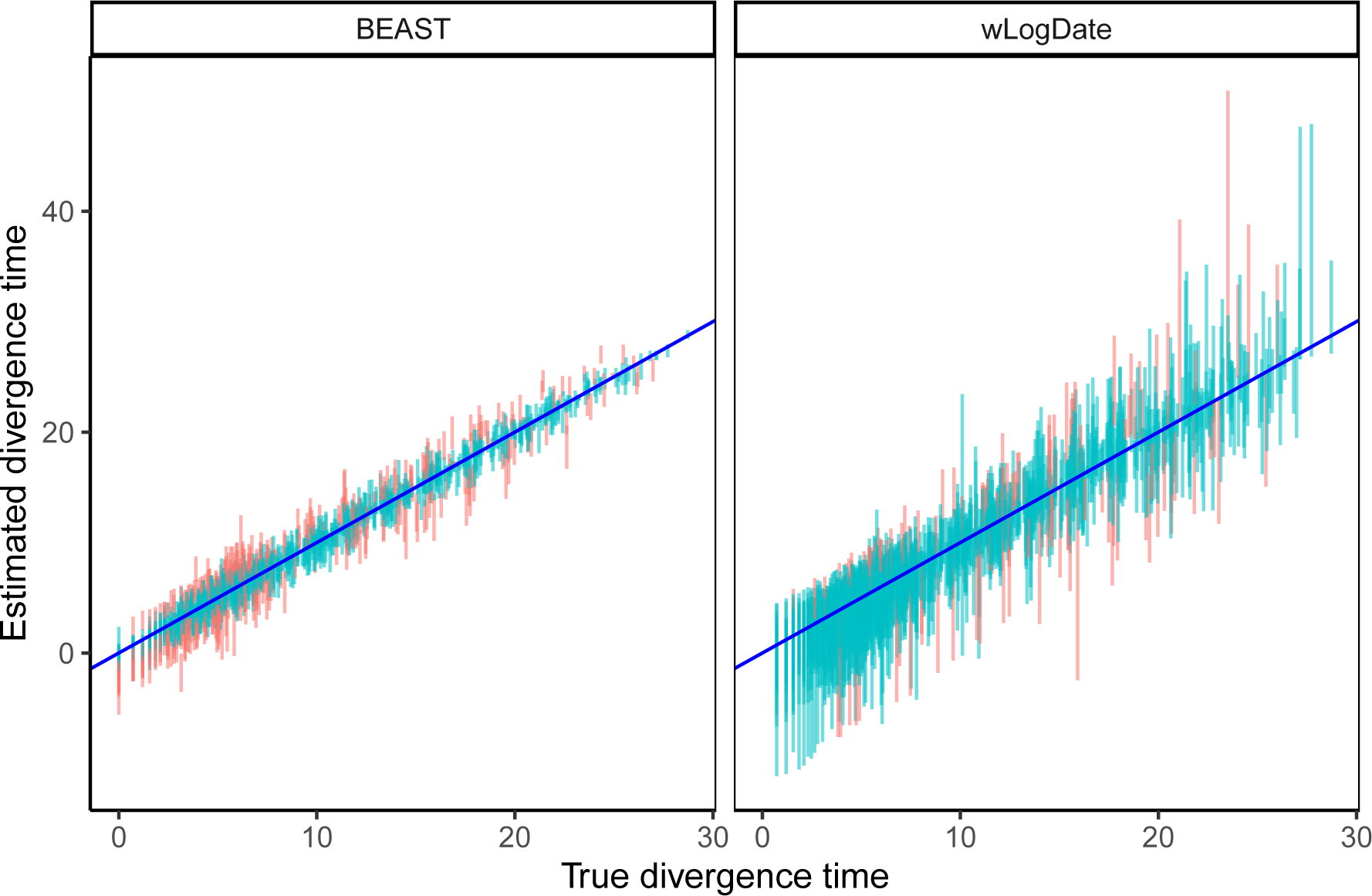
Estimated versus true divergence time. Each bar corresponds to the 95% confidence interval (CI) of one node estimate (each of the 109 nodes of the 10 replicates) by BEAST strict clock and wLogDate. Red color is used to mark points where the true time falls outside the CI.

In terms of the mutation rate, the distinction between methods is less pronounced (Table S1). wLogDate is the best method jointly with the two strict clock models BEAST-strict and LF. Overall, even though LF and wLogDate tend to over-estimate mutation rates, both have less biased results compared to other methods (Fig. 4). LSD and BEAST-LogNormal have the highest errors; depending on the condition, each can overestimate or underestimate the rate but LSD tends to underestimate while BEAST-LogNormal tends to overestimate. On M1, wLogDate and LF have a clear advantage over BEAST-strict, which tends to over-estimate the rate. On M2, the three methods have similar accuracy. For M3 and M4, BEAST-strict under-estimates the rate under the Exponential model of rate variation, and wLogDate and LF are closer to the true value. For all methods, M4 is the most challenging case.

We also compare confidence intervals obtained from wLogDate and BEAST (Fig.5). Although wLogDate intervals are on average 2.7 times larger than BEAST, 33% and 12% of the true values fall outside the 95% confidence interval for BEAST and wLogDate, respectively. Thus, while both methods under-estimate the confidence interval range, wLogDate, with its larger intervals, is closer to capturing the true value in its confidence interval at the desired level.

Finally, we compared all methods in terms of their running time (Table S2). LSD and LF are the fastest methods in all conditions, always taking tens of seconds (less than a minute) on these data. The running time of wLogDate depends on the model condition and can be an order of magnitude higher for Exponential rates than the other two models of rate variation.

Nevertheless, wLogDate finishes on average in half a minute to 12 minutes, depending on the model condition. In contrast, BEAST took close to one hour with strict clock and close to two hours with the LogNormal model (and even more if run with longer chains; see Table S5 in Supplementary Materials.

### 3.2 Simulated data with autocorrelated rate

In simulations with the autocorrelated model, we compare LF, RelTime, and wLogDate (Fig. 6). The distribution of the estimated divergence time of uncalibrated internal nodes does not show any sign of biased in divergence time estimation for either method. All methods seem to give less varied estimates for the younger nodes (i.e. those with higher divergence times) and have more varied estimates for older nodes. In addition, the estimates of wLogDate are more concentrated around the true values than that of LF and RelTime, indicating a better accuracy. In two test cases (out of 200), LF had extremely high error (Fig.S7). Once those two cases are removed, the average RMSE normalized by tree height is 0.09 for wLogDate, 0.10 for LF, and 0.13 for RelTime (Table S7). Comparing to LF and wLogDate, RelTime gives wider distributions of the estimates for a large portion of the nodes.

**Figure 6:**
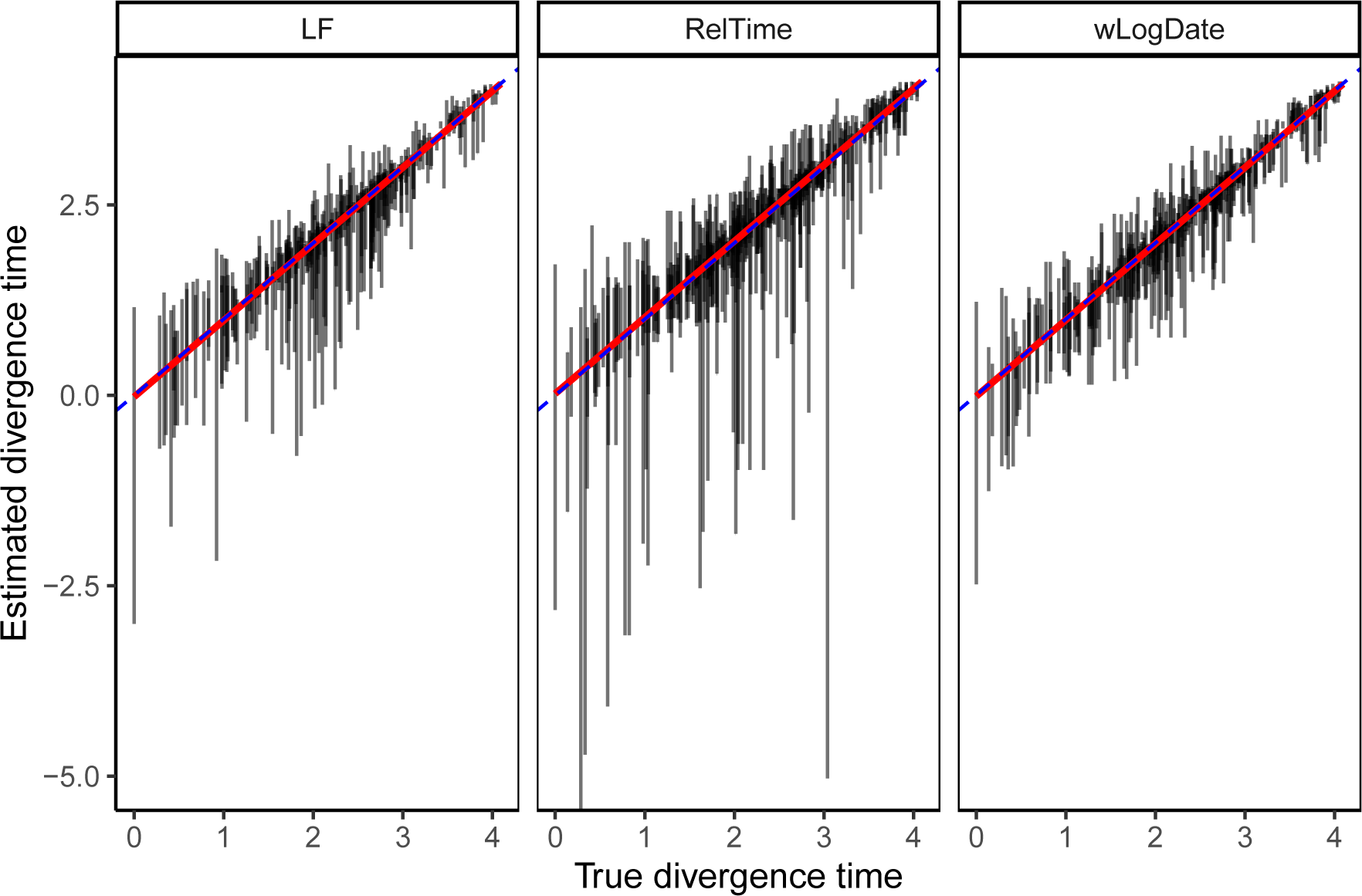
Comparison of LF, RelTime, and wLogDate on the simulated data with autocorrelated rate model. The y-axis shows estimated divergence times of uncalibrated internal nodes while the x-axis shows the true divergence time. Each bar shows the 2.5% and 97.5% quantiles of the estimates of a single node’s divergence time across 20 tests, each of them with different random choices of calibration points (thus, these are not CIs for one run). There are 10 replicate trees, each with 44 uncalibrated nodes (thus, 440 bars in total). This figure discards 2 tests (out of 10 × 20 = 200) where LF produced extremely erroneous time trees (see Fig. S7) for the full results). The root-mean-square error of the un-calibrated internal node ages, normalized by the tree height averaged across all replicates were 0.09, 0.1, and 0.13, respectively, for wLogDate, LF, and RelTime (see Table S7).

### 3.3 Biological data

On the H1N1 dataset, the best available evidence has suggested a tMRCA between December 2008 and January 2009 [15, 27, 34]. wLogDate inferred the tMRCA to be 14 December 2008 (Fig. 7a), which is consistent with the literature. LF and LSD both infer a slightly earlier tMRCA (10 November 2008), followed by BEAST-strict and BEAST-lognorm (October 2008 and July 2008), and finally BEAST runs using the phyML tree (Feb. 2008 for strict and July 2007 for LogNormal). While the exact tMRCA is not known on this real data, the results demonstrate that wLogDate, on a real data, produces times that match the presumed ground truth.

**Figure 7:**
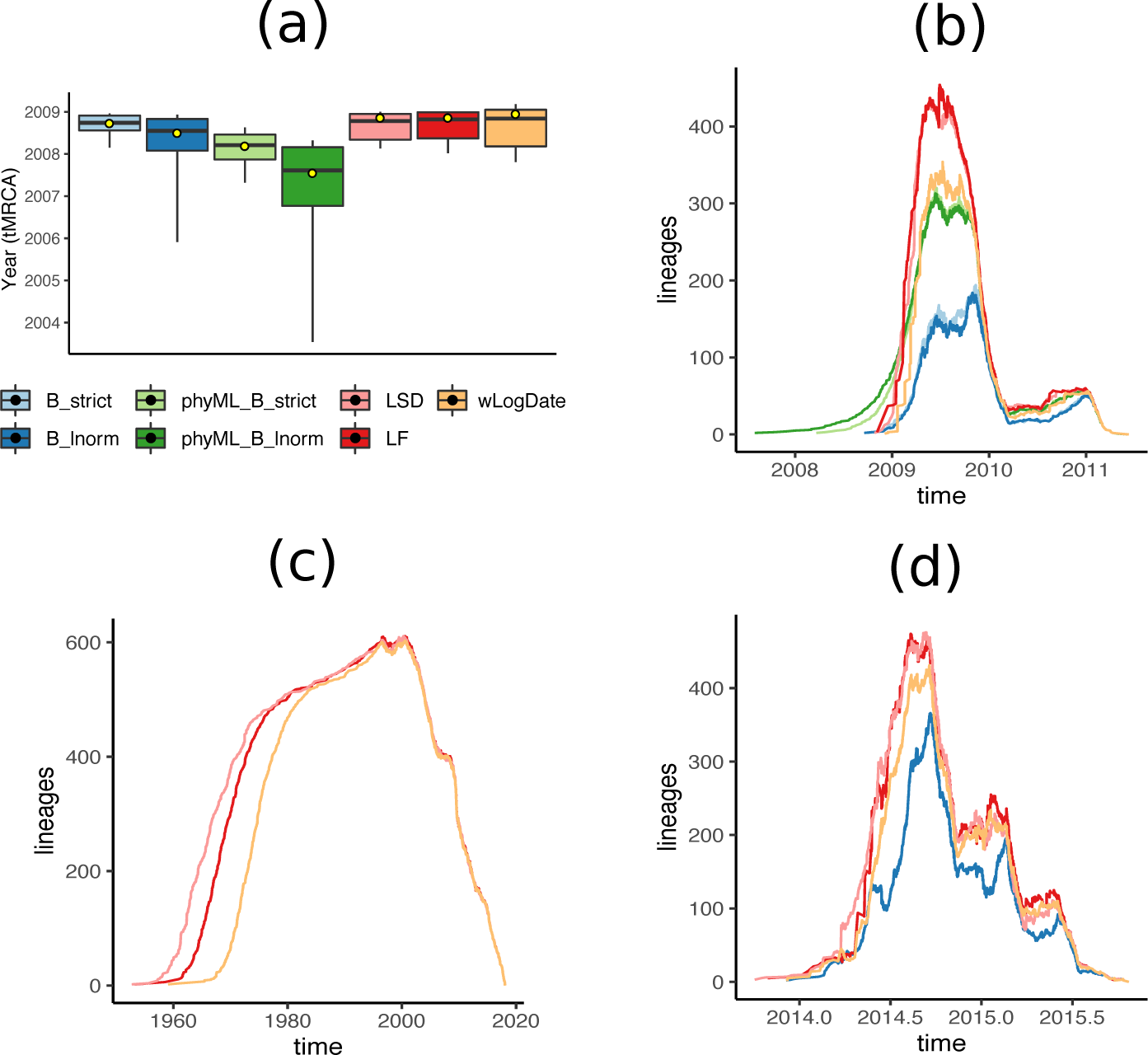
(a) Inferred tMRCA of the H1N1 dataset. Boxplots represent the median, maximum, minimum, 97.5% and 2.5% quantiles of the bootstrap estimates for LF, LSD, and wLogDate, and of the posterior distribution for BEAST. Yellow dot shows the inferred tMRCA of the best ML or MAP tree. BEAST was run with 4 different settings: B strict and B lnorm allow BEAST to infer both tree topology and branch lengths, with strict and LogNormal clock models; phyML B strict and phyML B lnorm fixed the topology to the rooted phyML tree given to BEAST. All other methods (LSD, LF, and wLogDate) were run on the rooted phyML trees. Results for LSD, LF, and BEAST are all obtained from To *et al*. [50]. (b) LTT plot for all methods on the H1N1 data. (c) LTT plot of fast methods on the HIV dataset. (d) LTT plot of BEAST, LSD, LF, and wLogDate on the Ebola dataset.

On the HIV dataset, wLogDate inferred a tMRCA of 1958 with only a handful of lineages coalescing in the 1950s and most others coalescing in 1960s and early 1970s (Fig. S3). The recovered tMRCAs is within the range postulated in the literature for subtype B [12, 53] and the fact that randomly sampled HIV lineages across USA tend to coalesce deep in the tree is a known phenomenon. LF and LSD recovered the tMRCA of 1952 and 1953, respectively. Comparing to wLogDate, these two strict-clock methods postulate an earlier burst of subtype B (Fig. 7c). We were not able to run BEAST on this dataset.

On the Ebola dataset, the BEAST-1000 trees obtained from Dudas *et al*. [7] inferred the tMRCA to be between 13 September 2013 and 26 January 2014 (95% credible interval) and the BEAST-MCC inferred the tMRCA to be 5 December 2013 as reported by Volz and Frost [51]. Here, wLogDate inferred a tMRCA on 7 December 2013, which is very close to the estimate by BEAST. Both LF and LSD inferred an earlier tMRCA: 29 October 2013 for LF and 2 October 2013 for LSD, but still within the 95 per cent credible interval of BEAST-1000. LTT plots showed a similar reconstruction by all methods for this dataset (Fig. 7d).

We also compare running times of dating methods on the three real biological datasets (Table S3). LSD was always the fastest, running in just seconds, compared to minutes for LF and wLogDate. LF is faster than wLogDate on the H1N1 and HIV data, while on Ebola data, wLogDate is faster. We report the running time for wLogDate as the sequential run of 10 independent starting points and note that wLogDate can easily be parallelized. We further tested the scaling of wLogDate with respect to the number of species by subsampling the HIV dataset to smaller numbers of species (Fig. S2). The results show that the running time of wLogDate increases slightly worse than quadratically with the incrased number of species.

## 4 Discussion and future work

We introduced (w)LogDate, a new method for dating phylogenies based on a non-convex optimization problem. We showed that by log-transforming the rates before minimizing their variance, we obtain a method that performs much better than LSD, which is a similar method without the log transformation. In phylodynamics settings, our relatively simple method also outperformed other existing methods, including the Bayesian methods, which are much slower. The improvements were most pronounced in terms of the estimation of tMRCA and individual node ages and less so for the mutation rate. Moreover, improvements are most visible under the hardest model conditions, and are also observed in when data are generated according to autocorrelated model of rates.

The log transformation results in a non-convex optimization problem, which is harder to solve than the convex problems solved by LSD and LF. However, we note that the problem is convex for rate multipliers between 0 and *e*. In addition, given the advances in numerical methods for solving non-convex optimization problems, insistence on convex problems seems unnecessary. Our results indicate that this non-convex problem can be solved efficiently in the varied settings we tested. The main benefits of the log transformation are two: (1) it allow us to define a scoring function that assigns symmetrical penalties for increased or decreased rates (Fig. 1a), and (2) it allows wider deviations from a strict clock.

The accuracy of LogDate under varied conditions we tested is remarkable, especially given its lack of reliance on a particular model of rate evolution. We emphasize that the parametric models used in practice are employed for mathematical convenience and not because of a strong biological reason to believe that they capture real variations in rates.

Even assuming biological realism of the rate model, the performance of the relaxed clock model used in BEAST was surprisingly low. For example, when rates are drawn from the LogNormal distribution, BEAST-strict often outperformed BEAST-LogNormal, especially in terms of the estimates of tMRCA and the mutation rate. We confirmed that the lower accuracy was not due to lack of convergence in the MCMC runs. We reran all experiments with longer chains (Table S5). to ensure ESS values are above 300 (Table S6). These much longer runs failed to improve the accuracy of the BEAST-LogNormal substantially and left the ranking of the methods unchanged (Fig. S8).

The LogDate approach can be further improved in several aspects. First, the current formulation of LogDate assumes a rooted phylogenetic tree, whereas most inferred trees are unrooted. Rooting phylogenies is a non-trivial problem and can also be done based on principles of minimizing rate variation [29]. Similar to LSD, LogDate can be generalized to unrooted trees by rooting the tree on each branch, solving the optimization problem for each root, and choosing the root that minimizes the (w)LogDate objective function. We leave the careful study of such an approach to the future work.

Beyond rooting, the future work can explore the possibility of building a specialized solver for LogDate to gain speedup. One approach could be exploiting the special structure of the search space defined by the tree, which is the strategy employed by LSD to solve the least-squares optimization in linear time. Divide-and-conquer may also prove effective.

The weighting scheme used in LogDate is chosen heuristically to deal with the deviations of estimated branch lengths from the true branch length. In future, the weighting schema should be studied more carefully, both in terms of theoretical properties and empirical performance.

We described, implemented, and tested LogDate in the condition where calibrations are given as exact times (for any combinations of leaves and internal nodes). While the current settings fit well to phylodynamics data, its application to paleontological data with fossil calibrations is somewhat limited due to the requirements for exact time calibrations (in contrast to the ability to allow minimum or maximum constraints on the ages, or a prior about the distribution of the ages as in BEAST and RelTime). While the mathematical formulation extends easily, treatment of fossil calibrations and uncertainty of times is a complex topic [14, 18] that requires the expansion of the current work. Applying LogDate for deep phylogenies would need further tweaks to the algorithm, including changing equality to inequality constraints and the ability to setup feasible starting points for the solver.

In the studies of LogDate accuracy, we have explored various models for rate heterogeinety, including parametric models where rates are drawn i.i.d. from a fixed distribution (Log-normal, Exponential, and Gamma) and autocorrelated model where the rates of adjacent branches are correlated. Overall, none of the methods we studied is the best under all conditions. In phylodynamics data, our simulations showed that it is more challenging for all the dating methods to date the phylogenies of the inter-host evolution (M3 and M4) than the intra-host (M1 and M2). wLogDate outperforms other methods for the inter-host phylogenies, regardless of the model of rate heterogeneity. While all methods have lower error for intra-host trees, BEAST with strict-clock prior is often the best method. However, the differences between BEAST and wLogDate are small and wLogDate is often the second best. Thus, wLogDate works well for virus phylogenies, especially in inter-host conditions. Despite the fact that RelTime explicitly optimizes the rate for each pairs of sister lineages, wLogDate is more accurate than both LF and RelTime on the data where the rates are autocorrelated between adjacent branches. These results show that wLogDate is applicable to a fairly large number of models of the trees and the rates.

Nevertheless, the approach taken by wLogDate suffers from its own limitations. By including a single mean rate around which (wide) variations are allowed, wLogDate is expected to work the best when rates have distribution that are close to being unimodal. However, rates on real phylogenies may have sudden changes leading to bimodal (or multimodal) rate distributions with wide gaps in between modes. For example, certain clades in the tree may have local clocks that are very different from other clades. Such a condition has been studied by Beaulieu *et al*. [2] for a dataset of seed plants. The authors setup a simulation where there are local clocks on the tree and the mean values of these clocks are different by a factor varying from 3 to 6. Beaulieu *et al*. [2] point out that under such condition, especially when the rate shift occurs near the root, BEAST usually overestimates the time of the Angiosperm (i.e. gives older time) by a factor of 2 (BEAST results from Beaulieu *et al*. [2] are reproduced in Fig. S9). We also tested wLogDate, LF, and RelTime on this dataset (Fig. S9). In scenario 2 of the simulation, where the rate shift between the two local clocks is extreme (a factor of 6), wLogDate clearly over-estimate the age of Angiosperms (by a median of 55%). In this same scenario, RelTime slightly underestimate the age (by 5%). In the other 4 scenarios where the rate shifts are more gentle, wLogDate continue to overestimate the age but by small margins (by 6%, 1%, 2%, and 5%), while RelTime underestimates ages also by small margins (3%, 5%, 4%, 3%, and 3%). LF has similar patterns to wLogDate. These results point to a limitation of wLogDate (and the other dating methods) in phylogenies with multiple local clocks.

In addition to multiple clocks, future works should test LogDate under models where rate continuously change with time, and have a direction of change. Finally, to facilitate the comparison between different methods, we used the true topology with estimated branch lengths. Future work should also study the impact of the incorrect topology on LogDate and other dating methods.

## Code availability

The LogDate code is available on https://github.com/uym2/LogDate.

## Data availability

All the data are available on https://github.com/uym2/LogDate-paper.

## 5 Acknowledgments

We thank Susan B. Little for providing the San Diego HIV sequence dataset used in this study. This work was supported by the National Science Foundation (NSF) grant III-1845967 and a developmental grant from the University of California, San Diego Center for AIDS Research (P30 AI036214), supported by the National Institutes of Health.

## Supplementary material

### 6 Supplementary text

#### Algorithm 1 Setup the linear constraints Ψ. for tree T given a set of calibration points.

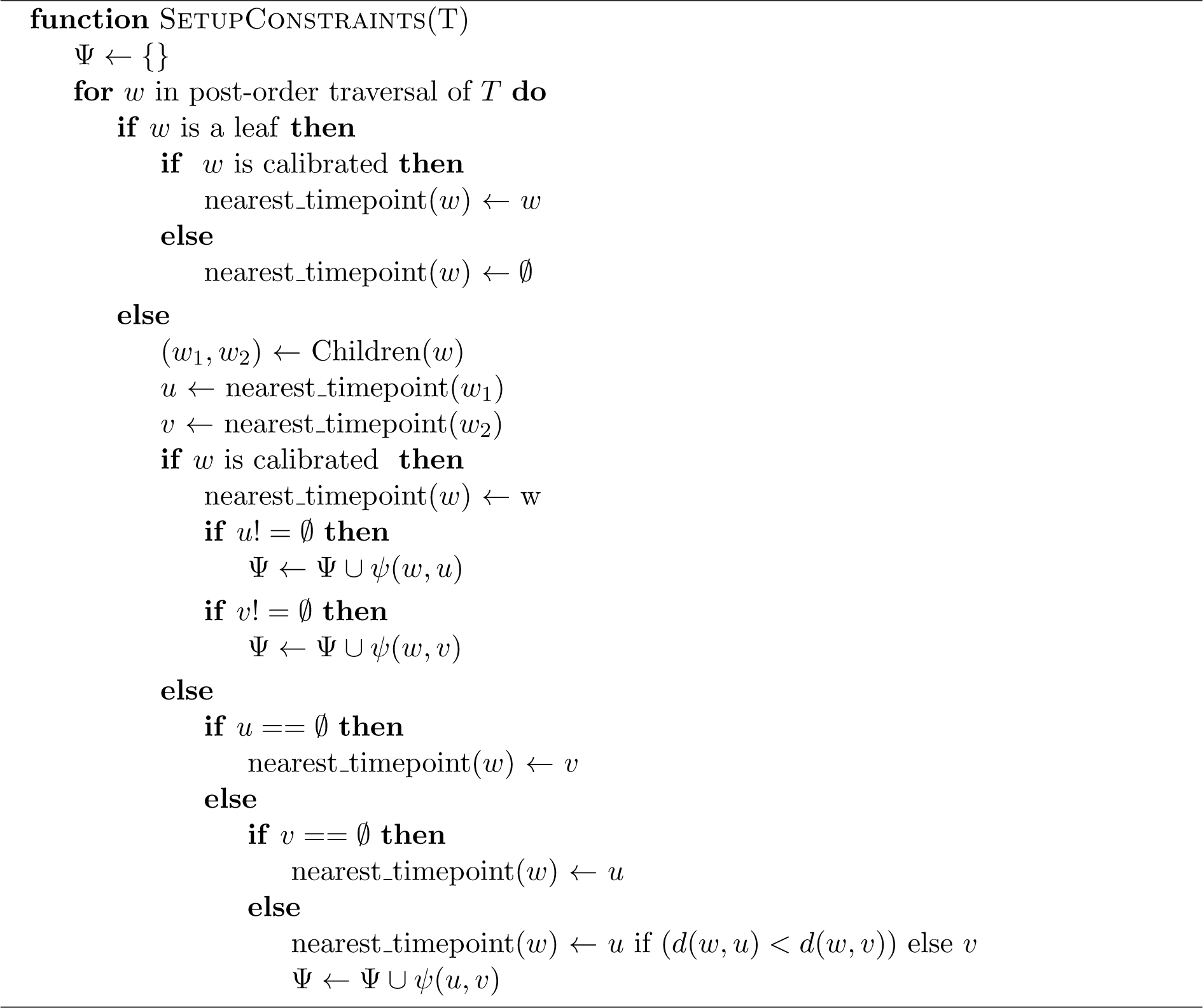

**Claim 1**. *The optimal* **x**^*^ *in Eq. 5 yields a set ν* = {*ν*_1_, *ν*_2_,…, *ν*_2*n*−1_} *that has the maximum joint probability under a model of rate variation where ν*_*i*_ *are i*·*i*·*d, ν*_*i*_ ∼ *LogNormal*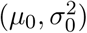,*and Mode*(*ν*_*i*_) = 1 *for any µ*_0_ *and* 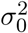, *subject to the constraints* Ψ.

*Proof*. We have: *v_i_* ~ *LogNormal* 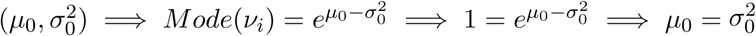. In other words, for the conditions with mode 1, we only have one free parameter.

The logarithm of the joint probability of *ν* under the LogNormal model of rate variation can be written as follows:

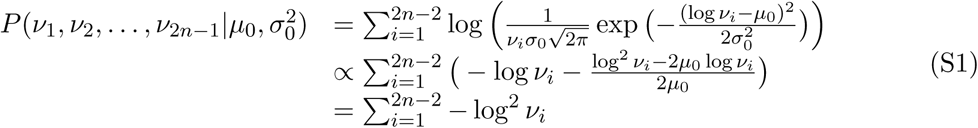

Thus, maximizing the joint probability of *ν* is equivalent to minimizing Eq. 5, subject to the constraints Ψ.

**Lemma 2**. *The length of the shortest path from the root of a binary tree to its leaves is at most* log *n where n is the number of leaves in the tree*.

*Proof*. Consider a rooted binary tree 𝒯 with *n* leaves; let *r* be the root and *h* be the length of the shortest path from *r* to the leaves of 𝒯. We need to prove that *h* ≤ log_2_ *n*.

Let 𝒟_*i*_ be the set of nodes in 𝒯 with depth *i*, that is, 𝒟_*i*_ = {*w* ∈ 𝒯 |*d*(*r, w*) = *i*}. We first prove that |𝒟_*i*_| = 2^*i*^ ∀*i* ≤ *h* where |𝒟_*i*_| denotes the cardinality of 𝒟_*i*_. We prove this by induction. The base case *i* = 0 holds since the root *r* is the only node with depth 0. Suppose we have |𝒟_*k*_| = 2^*k*^ and *k* < *h*, we need to prove that if *k* + 1 ≤ *h* then |𝒟_*k*+1_| = 2^*k*+1^. Note that a node *v* ∈ 𝒟_*k*+1_ if and only if its parent *par*(*v*) ∈ 𝒟_*k*_. Because 𝒯 is a binary tree, each node in 𝒯 must either has no child (leaf node) or two children (internal node). Since *k* < *h*, there must be no leaf node in 𝒟_*k*_, otherwise, a leaf *v* in 𝒟_*k*_ has *d*(*r, v*) = *k* < *h*, which defines a root-to-leaf path that is shorter than *h* and contradicts the definition of *h*. Thus, each node in 𝒟_*k*_ has exactly 2 children, making |𝒟_*k*+1_| = 2 * |𝒟_*k*_| = 2 * 2^*k*^ = 2^*k*+1^.

Now we have |𝒟_*h*_| = 2^*h*^. To prove that *h* ≤ log_2_ *n*, note that 𝒟_*h*_ contains a mixture of leaves and internal nodes and each internal node in 𝒟_*h*_ must have more than one leaf below it. Therefore, the size of 𝒟_*h*_ is at most the size of the leaf set of 𝒯; that is, |𝒟_*h*_| ≤ *n*. Thus, we have 2^*h*^ = |𝒟_*h*_| ≤ *n* =⇒ *h* ≤ log_2_ *n*.

**Claim 3**. *If all the leaves have sampling times and there is no other calibration points given for internal nodes, the matrix corresponding to the constraints* Ψ *setup by Algorithm 1 has O*(*n* log(*n*)) *non-zero elements, where n is the number of leaves in the input tree* 𝒯.

*Proof*. Let *P*(*w*) denote the shortest path from a node *w* to its leaves and let |*P*(*w*)| denote the length of this path. Let 𝒯_*w*_ be the clade of 𝒯 below *w* and let |*w*| denote the size of this clade (i.e. the number of leaves below *w*). Applying lemma 2 on 𝒯_*w*_, we have |*P*(*w*)| ≤ log_2_ |*w*| ≤ log_2_ *n* for all *w* ∈ 𝒯.

Note that if all leaves have sampling times, Algorithm 1 adds exactly one constraint for each internal node in the tree. For each node *w* with two children *c*_*l*_(*w*) and *c*_*r*_(*w*), the non-zero elements of the constraint added when node *w* is visited must locate on *P*(*c*_*l*_(*w*)), *P*(*c*_*r*_(*w*)), and the two branches (*w, c*_*l*_(*w*)) and (*w, c*_*r*_(*w*)). Let *η*_*w*_ denote the number of non-zero elements of the constraint defined by node *w*, then *η*_*w*_ ≤ |*P*(*c*_*l*_(*w*))| + |*P*(*c*_*r*_(*w*))| + 1 + 1 ≤ 2 log_2_ *n* + 2. Thus, the total number of non-zeros in all constraints corresponding to the *n* − 1 internal nodes is bounded above by (*n* − 1)(2 log_2_ *n* + 2) ∈ *O*(*n* log *n*).

### 7 Hybrid rate Angiosperm

Beaulieu *et al*.([2]) simulated a hybrid rate model for a phylogeny of seed plants in which evolutionary rates formed local clocks in certain clades of the tree. The authors simulated that data in 5 scenarios where they change the relative ratios between some clades in the tree, as follow:

- scenario 1 = 3:1 herbaceous to woody
- scenario 2 = 6:1 herbaceous to woody
- scenario 3 = 4:1 angio. to gymno.; 3:1 herbaceous to woody angio.
- scenario 4 = 4:1 angio. to gymno.; 3:1 herbaceous to woody angio.; Gnetales herbaceous angio.
- scenario 5 = 4:1 angio. to gymno.; 3:1 herbaceous to woody angio.; Gnetales woody angio.

The time tree and 100 simulated phylograms for each of these five scenarios were downloaded from the Dryad Repository provided by the authors. We used the provided phylograms to estimate the time tree using wLogDate, RelTime, and LF and compare the estimated age of Angiosperm to the true tree. Without the simulated sequences, we could not run BEAST. However, we show the BEAST results reported by the original study ([2]). We aware that the comparison to BEAST must be made with cautions, because the experimental settings were different as we will state below:

- As RelTime cannot run without outgroups, we had to use the 20 species on the clade outside the Angiosperm as outgroups. As such, this entire clade is ignored by RelTime and only 91 species out of 111 are included in the time tree. We used the same setting for wLogDate and LF. Because of this fact, 5 calibration points belong to the 20 species in the outgroups are also discarded out of the total 20 calibration points. However, in their original study, the authors ran BEAST using 20 calibration points (instead of 15 points) to date the full tree with 111 species (instead of 91 species).
- In their original study, the authors gave BEAST a distribution instead of exact-time for each calibration, as opposed to the exact-time points as we used to run LF, RelTime, and wLogDate.

## Supplementary figures and tables

**Table S1:** Mean absolute error of the inferred mutation rate of BEAST strict, BEAST lognorm, LF, LSD, and wLogDate.

| Tree Model | Clock Model | BEAST_strict | BEAST_lnorm | LF | LSD | RTT | wLogDate |
| --- | --- | --- | --- | --- | --- | --- | --- |
| M1 | Lognormal | 0.0007 | 0.0011 | **0.0004** | 0.0005 | 0.0007 | **0.0004** |
|  | Gamma | 0.0009 | 0.0014 | 0.0005 | **0.0004** | 0.0009 | 0.0006 |
|  | Exponential | 0.0009 | 0.0022 | **0.0008** | 0.0013 | 0.0013 | 0.0009 |
| M2 | Lognormal | **0.0005** | 0.0014 | **0.0005** | **0.0005** | 0.0008 | **0.0005** |
|  | Gamma | **0.0006** | 0.0013 | 0.0007 | 0.0007 | 0.0009 | 0.0007 |
|  | Exponential | **0.0013** | 0.0038 | 0.0015 | 0.0020 | 0.0019 | 0.0015 |
| M3 | Lognormal | **0.0003** | **0.0003** | 0.0006 | 0.0004 | 0.0008 | 0.0006 |
|  | Gamma | **0.0003** | **0.0003** | 0.0006 | 0.0004 | 0.0006 | 0.0006 |
|  | Exponential | 0.0010 | 0.0011 | **0.0009** | 0.0027 | 0.0012 | **0.0009** |
| M4 | Lognormal | **0.0007** | 0.0008 | 0.0008 | **0.0007** | 0.0010 | 0.0008 |
|  | Gamma | **0.0006** | 0.0007 | 0.0008 | 0.0008 | 0.0010 | 0.0007 |
|  | Exponential | 0.0020 | **0.0016** | **0.0016** | 0.0037 | 0.0018 | 0.0017 |
| Average |  | **0.0008** | 0.0013 | **0.0008** | 0.0012 | 0.0011 | **0.0008** |

**Table S2:** Average running time (seconds) of BEAST strict, BEAST lognorm, PhyML + LF, PhyML + LSD, and PhyML + wLogDate with 10 initials on simulated data.

| Tree Model | Clock Model | BEAST_strict | BEAST_lnorm | LSD | LF | wLogDate |
| --- | --- | --- | --- | --- | --- | --- |
| M1 | Lognormal | 2784.01 | 6018.80 | 15.01 | 17.47 | 73.65 |
|  | Gamma | 2823.09 | 6082.62 | 17.81 | 20.29 | 83.19 |
|  | Exponential | 2696.61 | 5840.18 | 17.44 | 19.40 | 723.61 |
| M2 | Lognormal | 2425.83 | 5207.20 | 18.61 | 19.94 | 39.32 |
|  | Gamma | 2466.24 | 5303.63 | 20.12 | 21.47 | 81.14 |
|  | Exponential | 2385.73 | 5169.07 | 20.87 | 22.12 | 418.01 |
| M3 | Lognormal | 3848.01 | 8204.00 | 35.25 | 38.07 | 55.92 |
|  | Gamma | 3850.71 | 8211.09 | 38.71 | 41.55 | 65.51 |
|  | Exponential | 3826.12 | 6520.19 | 40.10 | 43.05 | 280.55 |
| M4 | Lognormal | 2914.03 | 6201.59 | 28.84 | 30.40 | 38.92 |
|  | Gamma | 2901.59 | 6184.01 | 30.78 | 32.29 | 41.56 |
|  | Exponential | 2855.79 | 6145.03 | 32.96 | 34.54 | 371.97 |

**Table S3:**
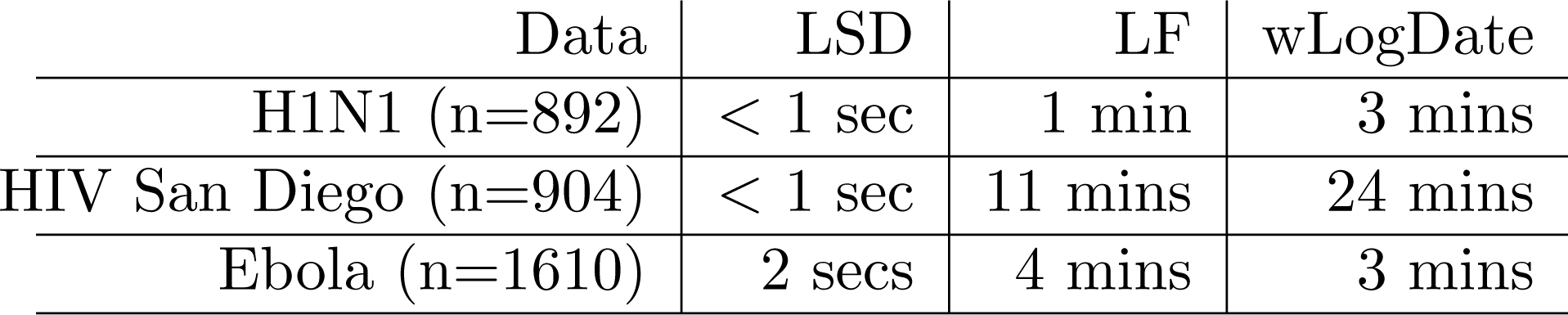
Running time of LSD, LF, and wLogDate on biological datasets.

**Table S4:** Average RMSE of the internal node ages inferred by different weight functions for LogDate. Numbers are rounded to the closest 3 decimal digits. Recall that 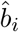 is the estimated branch length and 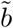 is a small constant.

| Tree Model | Clock Model | $w_i = 1$ | $\hat{b}_i + \tilde{b}$ | $(\hat{b}_i + \tilde{b})^2$ | $\log(1 + \hat{b}_i + \tilde{b})$ | $\sqrt{\hat{b}_i + \tilde{b}}$ | $\log(1 + \sqrt{\hat{b}_i + \tilde{b}})$ |
| --- | --- | --- | --- | --- | --- | --- | --- |
| M1 | Lognormal | 0.020 | 0.019 | 0.027 | 0.019 | **0.018** | **0.018** |
| M1 | Gamma | 0.020 | 0.018 | 0.026 | 0.018 | **0.017** | **0.017** |
| M1 | Exponential | 0.363 | 0.051 | 0.062 | 0.048 | **0.037** | 0.039 |
| M2 | Lognormal | 0.107 | 0.043 | 0.062 | 0.043 | **0.038** | 0.042 |
| M2 | Gamma | 0.100 | 0.046 | 0.064 | 0.046 | **0.043** | 0.044 |
| M2 | Exponential | 0.359 | 0.135 | 0.166 | 0.129 | **0.099** | 0.100 |
| M3 | Lognormal | **0.039** | 0.051 | 0.134 | 0.050 | **0.039** | **0.039** |
| M3 | Gamma | 0.041 | 0.050 | 0.132 | 0.049 | **0.039** | **0.039** |
| M3 | Exponential | 0.220 | 0.173 | 0.349 | 0.168 | 0.105 | **0.103** |
| M4 | Lognormal | 0.074 | 0.098 | 0.172 | 0.085 | **0.070** | 0.071 |
| M4 | Gamma | 0.098 | 0.086 | 0.172 | 0.082 | 0.070 | **0.069** |
| M4 | Exponential | 0.578 | 0.397 | 1.042 | 0.349 | 0.301 | **0.283** |

**Table S5:** BEAST convergence analysis: BEAST was run on Lognormal clock models with the correct prior. For each tree model, we run BEAST with a sufficiently long MCMC chain to ensure the effictive-sample-size (ESS) of all parameters are at least 200. We report the length of the MCMC chain, relative error of node age estimates with respect to BEAST using 10 millions MCMC chain, and the running time.

| Tree Model | MCMC Chain | Relative Error | Run Time (hours) |
| --- | --- | --- | --- |
| M1 | $10^7$ | 1.00 | 1.66 |
| M2 | $5 \times 10^7$ | 0.99 | 7.6 |
| M3 | $5 \times 10^7$ | 0.99 | 11.3 |
| M4 | $20 \times 10^7$ | 0.98 | 35.6 |

**Table S6:** BEAST convergence analysis: BEAST was run on Lognormal clock models with the correct prior. For each tree model, we run BEAST with a sufficiently long MCMC chain to ensure the effictive-sample-size (ESS) of all parameters are at least 200. We report the average ESS of posterior, likelihood, rootHeight, and meanRate of the first 10 replicates of each tree model.

| Tree Model | Posterior ESS | Likelihood ESS | MeanRate ESS | RootHeight ESS |
| --- | --- | --- | --- | --- |
| M1 | 420.5 | 753.3 | 382.0 | 406.9 |
| M2 | 766.9 | 4224.4 | 430.1 | 647.3 |
| M3 | 1242.3 | 4024.1 | 531.0 | 430.3 |
| M4 | 1254.9 | 16336.7 | 744.1 | 847.2 |

**Table S7:**
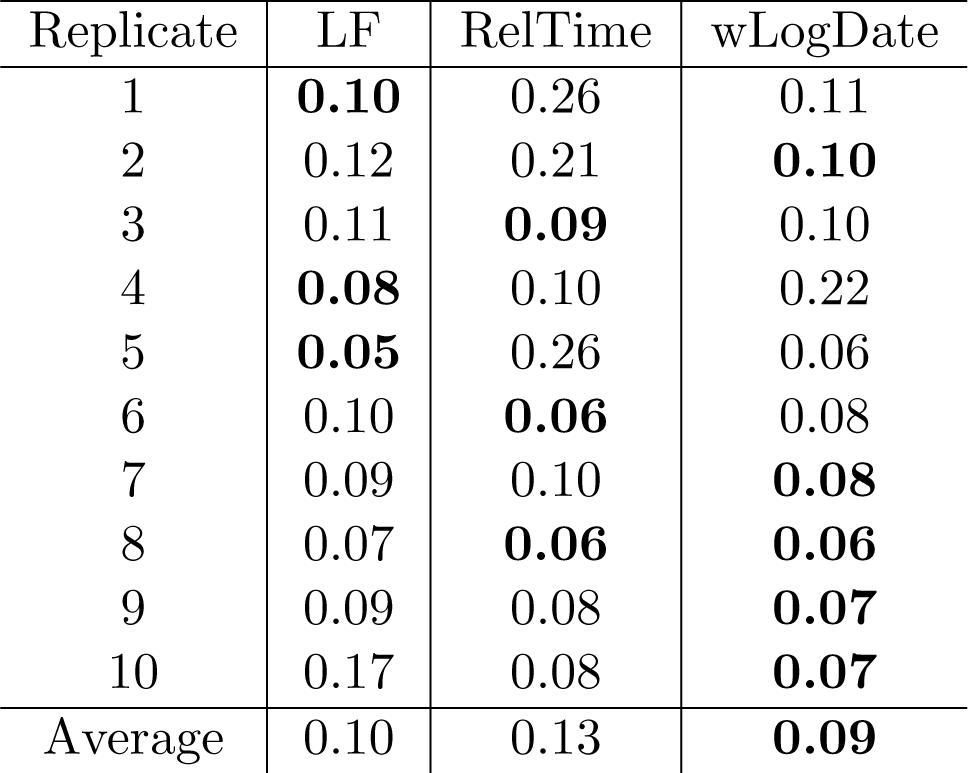
Comparison of LF, RelTime, and wLogDate on autocorrelated rate dataset. The Root-mean-square error (RMSE) of un-calibrated internal node ages is normalized by the tree height and reported for each replicate. Results discarded the two tests where LF produced extremely erroneous time tree. Refer to Fig. S7 for a complete picture.

| Replicate | LF | RelTime | wLogDate |
| --- | --- | --- | --- |
| 1 | **0.10** | 0.26 | 0.11 |
| 2 | 0.12 | 0.21 | **0.10** |
| 3 | 0.11 | **0.09** | 0.10 |
| 4 | **0.08** | 0.10 | 0.22 |
| 5 | **0.05** | 0.26 | 0.06 |
| 6 | 0.10 | **0.06** | 0.08 |
| 7 | 0.09 | 0.10 | **0.08** |
| 8 | 0.07 | **0.06** | **0.06** |
| 9 | 0.09 | 0.08 | **0.07** |
| 10 | 0.17 | 0.08 | **0.07** |
| Average | 0.10 | 0.13 | **0.09** |

**Figure S1:**
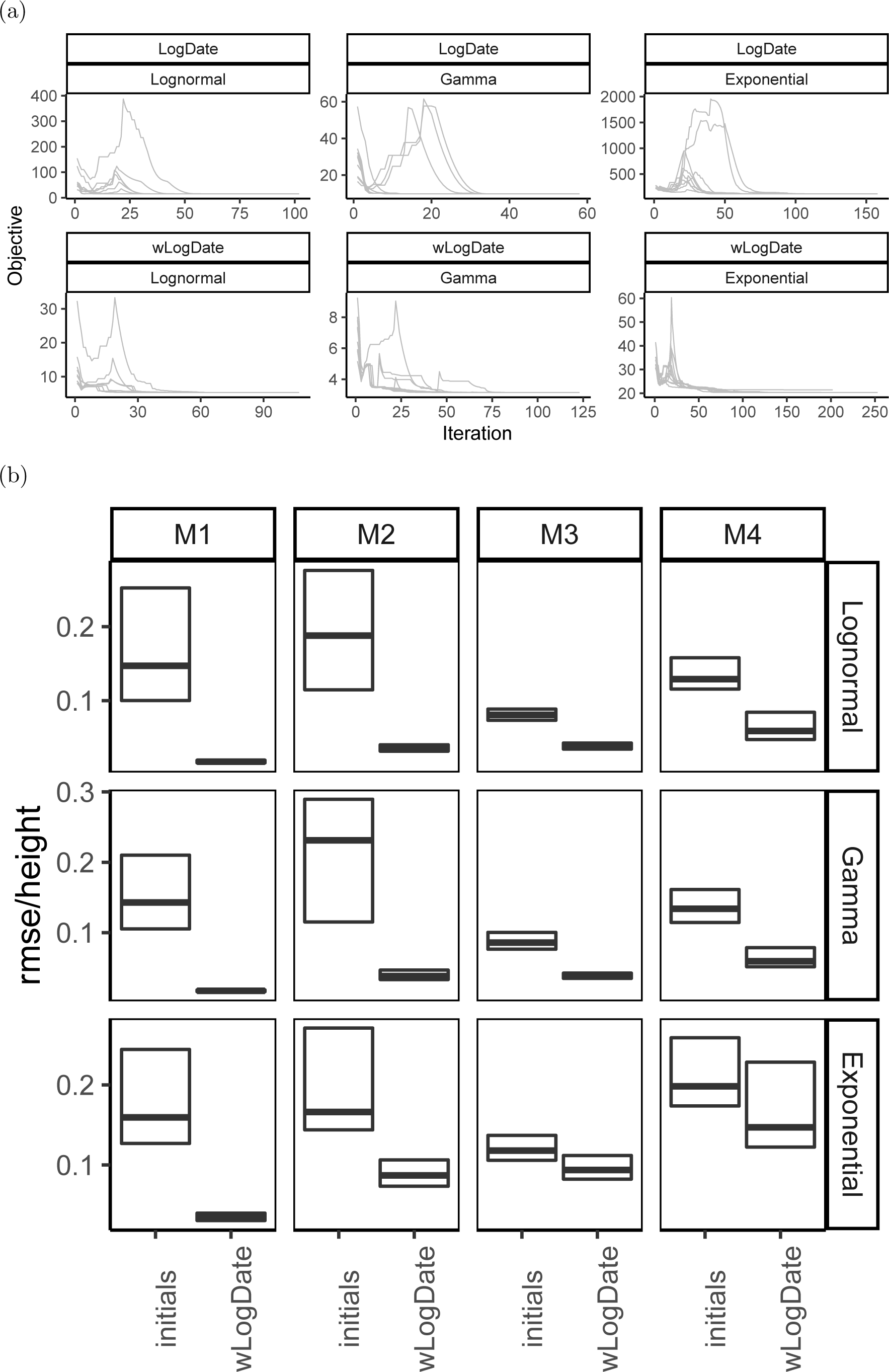
(a) Objective value versus iteration of the LogDate and wLogDate runs on one arbitrarily selected simulated tree (M4, replicate 2). Each of the two methods were run using 10 random initial points generated using the strategy described in the main text. (b) Normalized root-mean-square error of wLogDate versus the 10 initials used to run wLo2g9Date.

**Figure S2:**
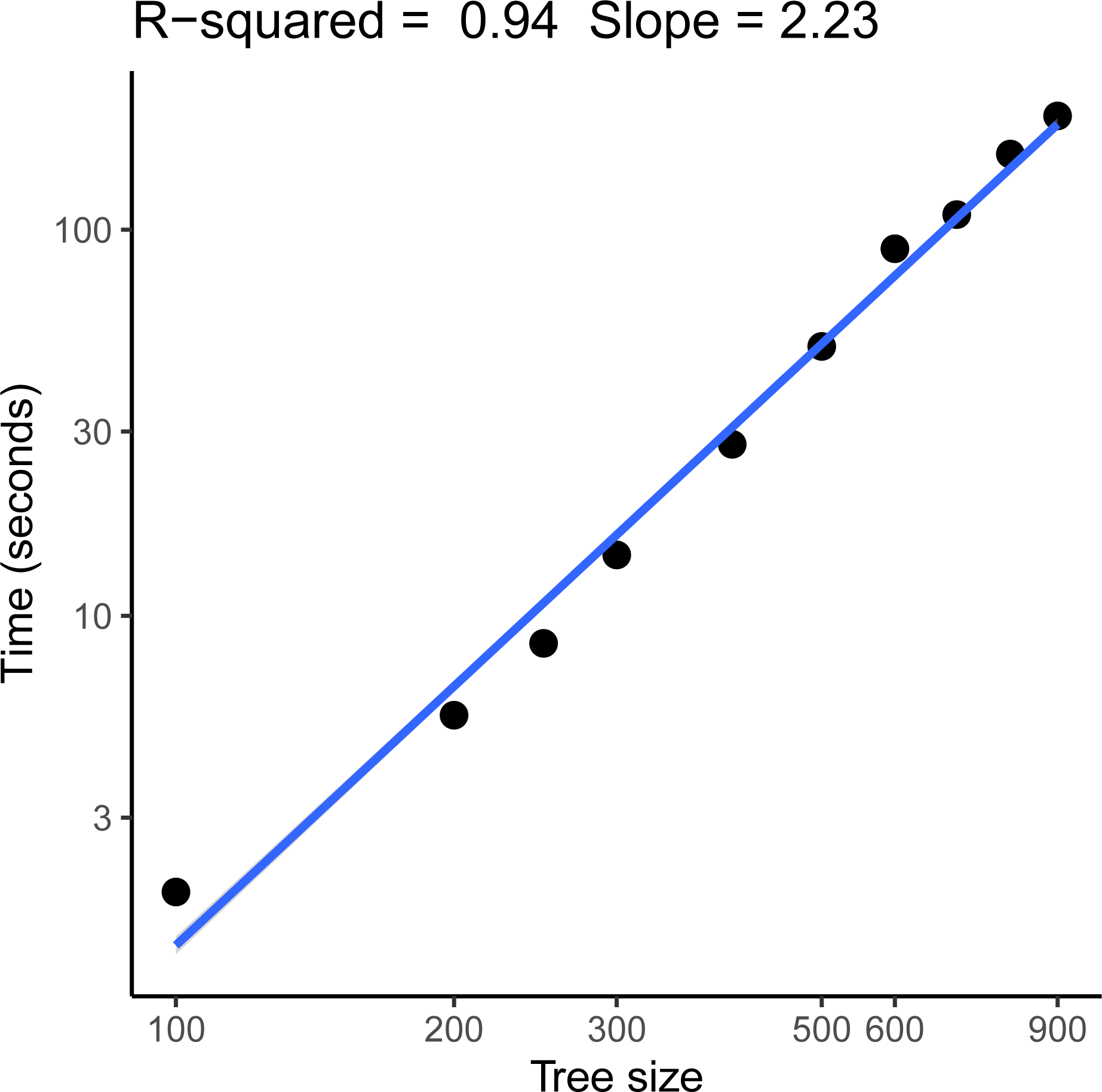
Running time of wLogDate on random subsets of the HIV dataset. For each tree size, wLogDate was run 100 times on 10 random subsets each with 10 initial points. Each dot represents the average run time of wLogDate per subset per initial point. Both axes are scaled in log (base 10). The slope of the line (2.23) shows the polynomial degree of the running time increase of wLogDate. Thus, wLogDate scales slightly worse than quadratically with increased numbers of species.

**Figure S3:**
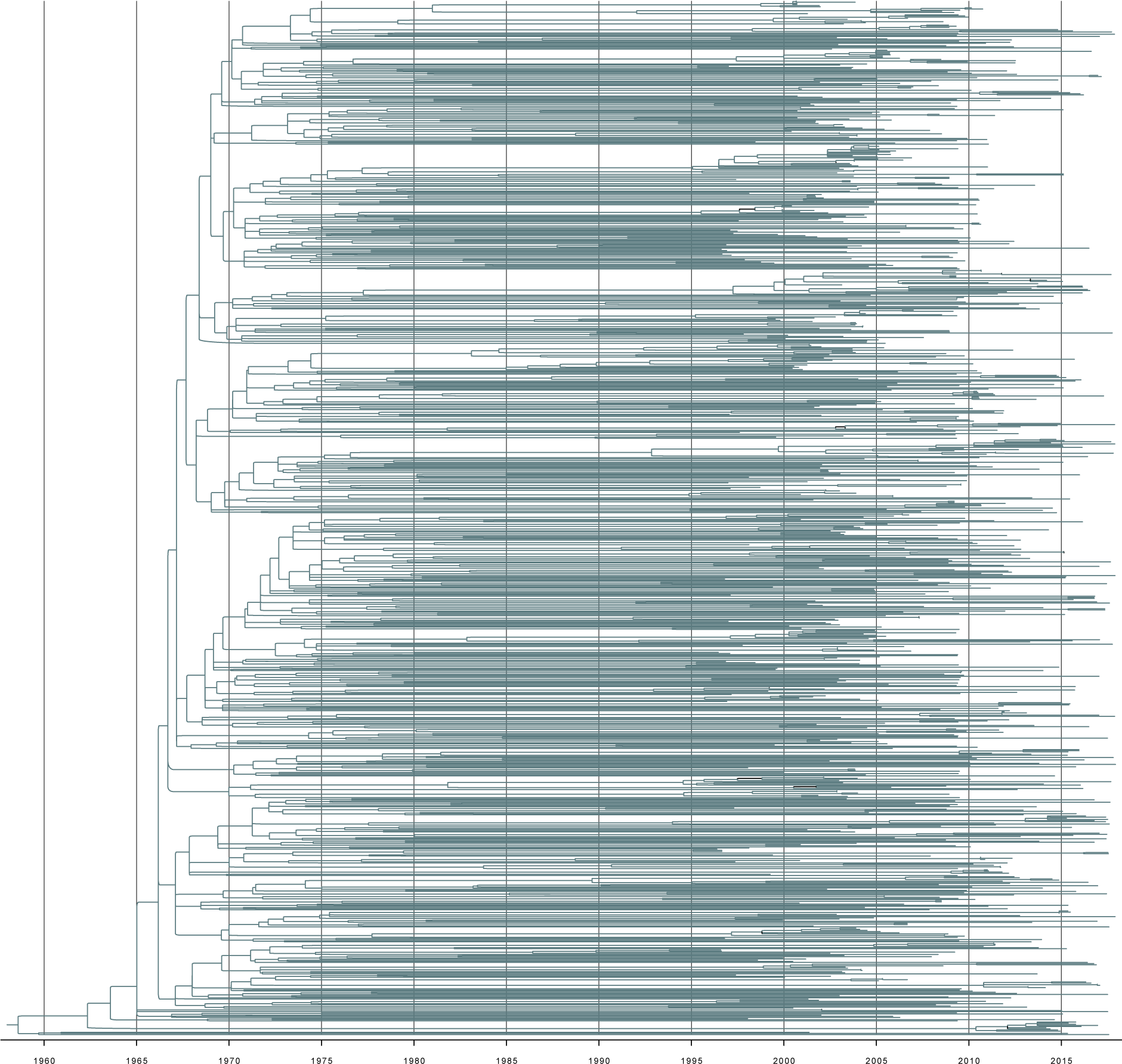
A TimeTree of San Diego HIV epidemic according to wLogDate.

**Figure S4:**
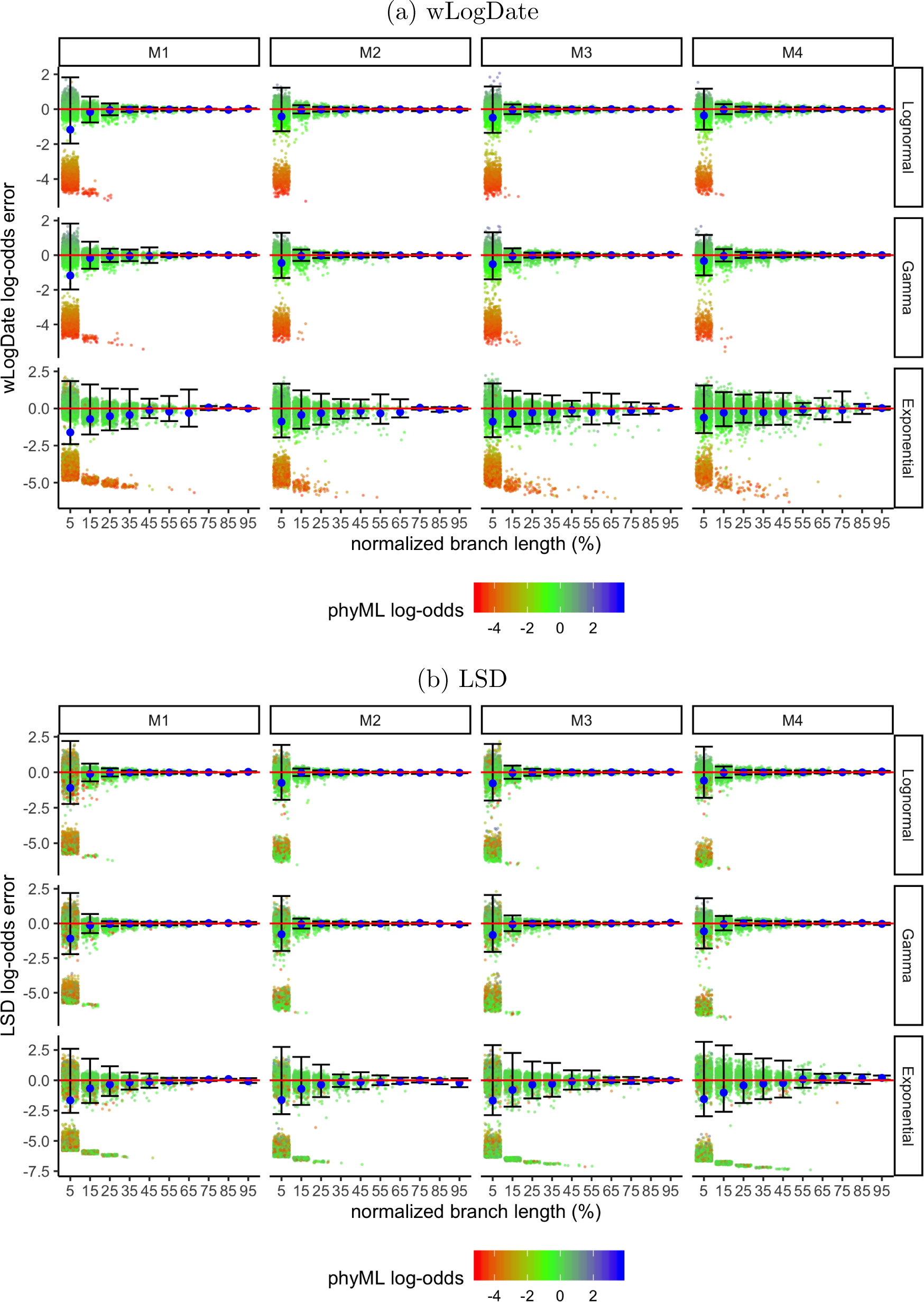
Effects of PhyML estimation error on wLogDate and LSD performance. Figure shows log-odds error of (a) wLogDate and (b) LSD versus true branch length (in time unit); x-axis is normalized by the maximum tree branch; dots are colored by log-odds error of phyML estimates; large blue dots show means and bars show one standard deviations around medians.

**Figure S5:**
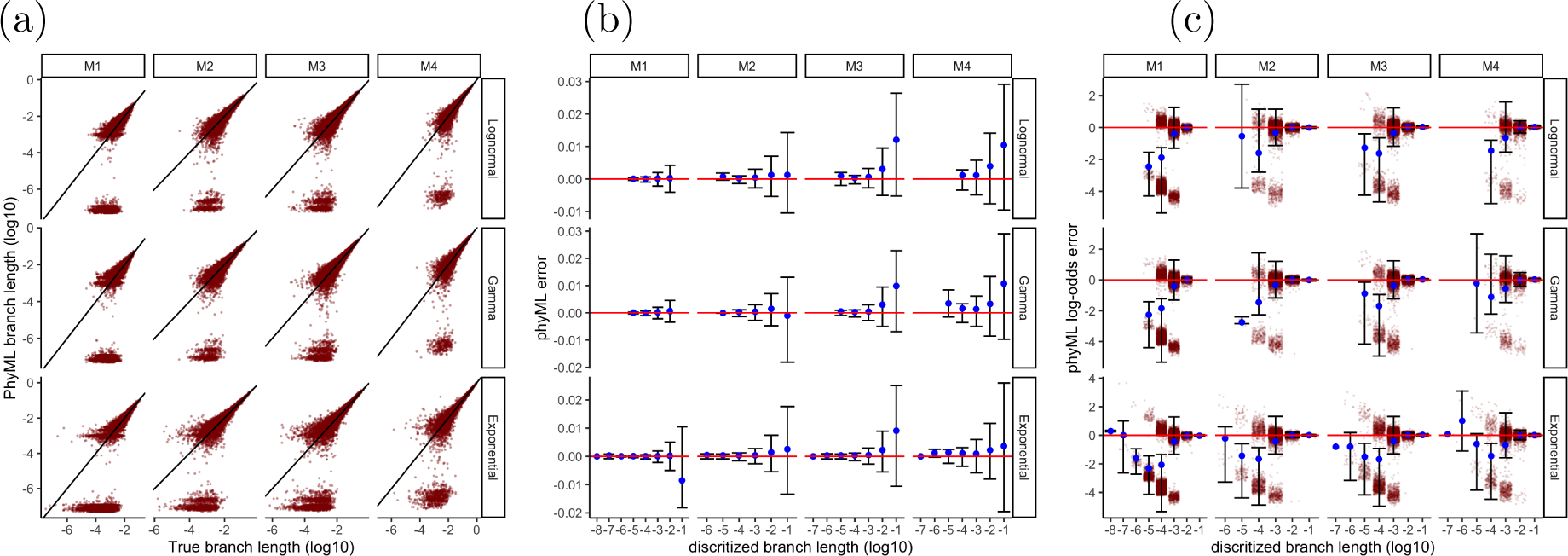
Analyses of the estimated branch lengths using PhyML on simulated data. (a) Estimated versus true branch lengths (expected number of substitutions per site); axes scaled in log10. (b) Error versus true branch lengths; blue dots represent means and bars represent standard deviations around medians. (c) Log-odds error versus true branch lengths; blue dots represent means and bars represent standard deviations around medians.

**Figure S6:**
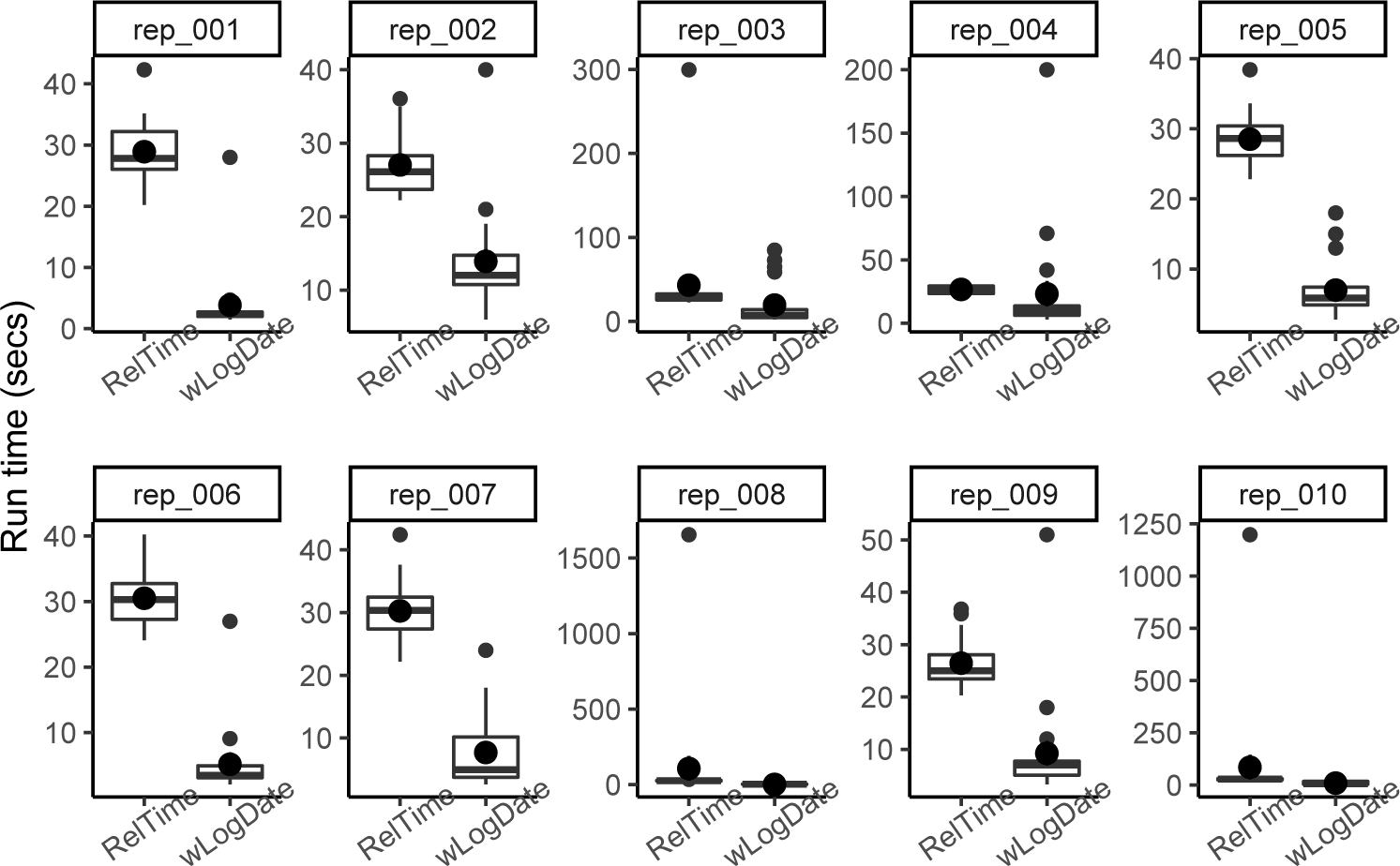
Run time of wLogDate and RelTime on the 10 replicates. Box plots show distributions for the 20 tests.

**Figure S7:**
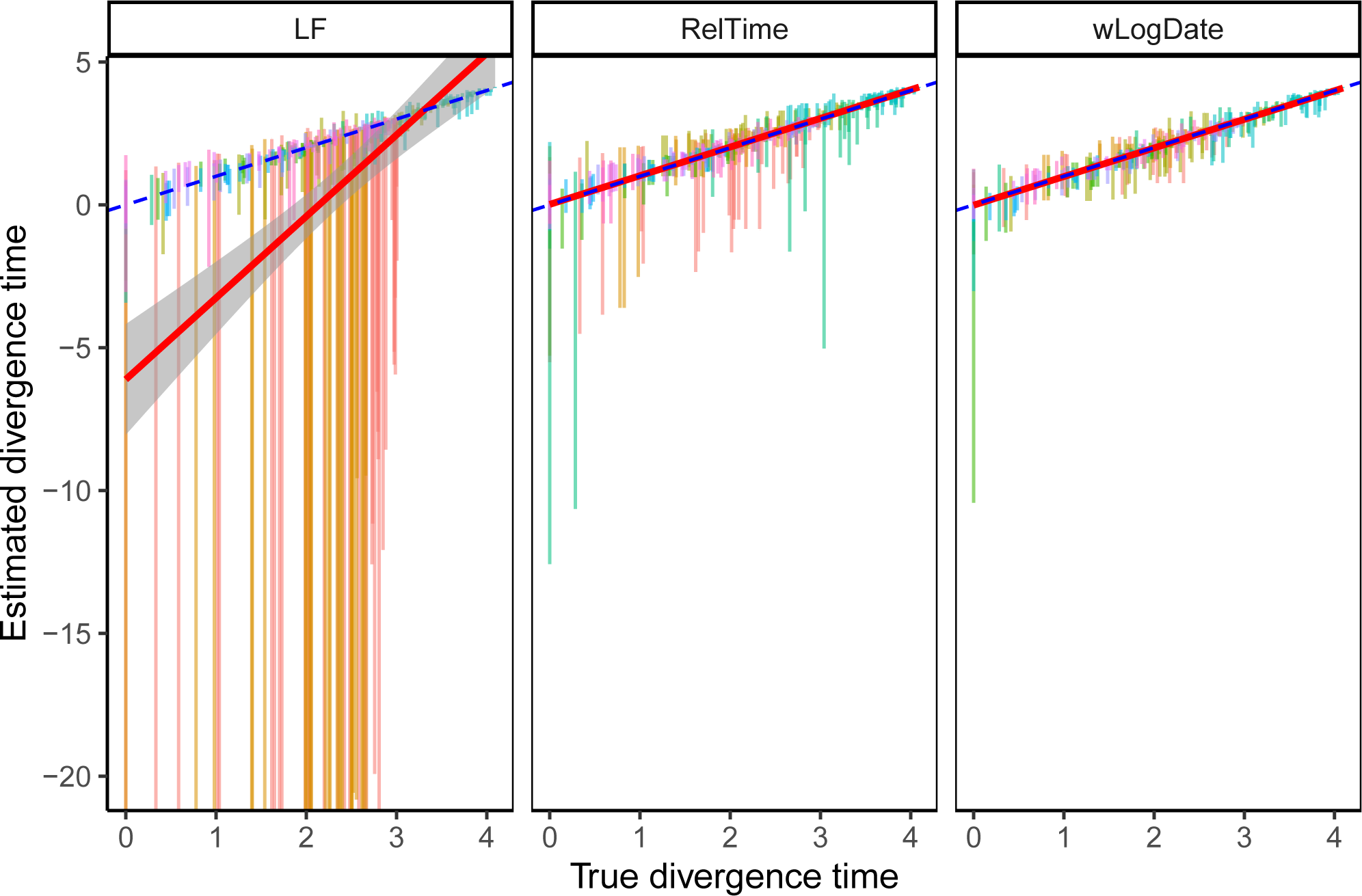
Comparison of LF, RelTime, and wLogDate on the simulated data with autocorrelated rate model. The y-axis shows estimated divergence times of uncalibrated internal nodes while the x-axis shows the true divergence time. Each bar shows the 2.5% and 97.5% quantiles of the estimates of a single node’s divergence time across 20 tests, each of them with different random choices of calibration points (thus, these are not CIs for one run). There are 10 replicate trees, each with 44 uncalibrated nodes (thus, 440 bars in total). There are two tests where LF produces extremely erroneous time trees (test 2 of replicate 1 and test 16 of replicate 2) and were discarded in Fig. 6 in the main text. The normalized RMSE of LF are 41.3 and 167.8 for these two tests, while the overall error without these two tests is 0.09. Here we show the full results for completeness. Colors are used to distinguish between replicates.

**Figure S8:**
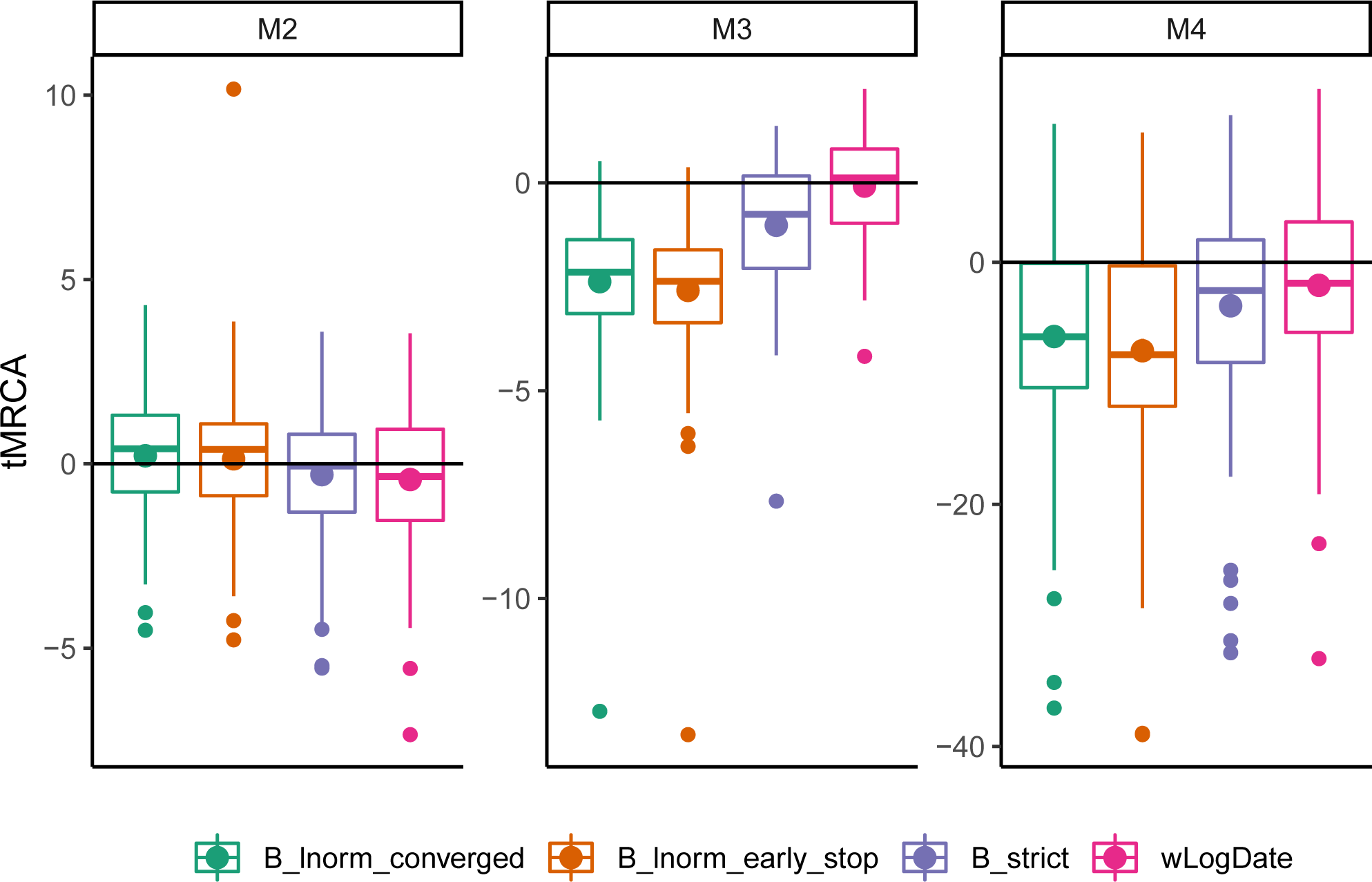
Estimation of the tMRCA of *M* 2, *M* 3, and *M* 4 of the simulated data with Lognormal clock model. For each model, BEAST was run with 3 conditions: B strict uses the strict-clock prior, B lnorm early stop uses Lognormal clock prior with MCMC chain of 10 millions, and B lnorm converged uses Lognormal clock prior with elongated MCMC chain to guarantee convergence (refer to table S6 for parameters and convergence check.)

**Figure S9:**
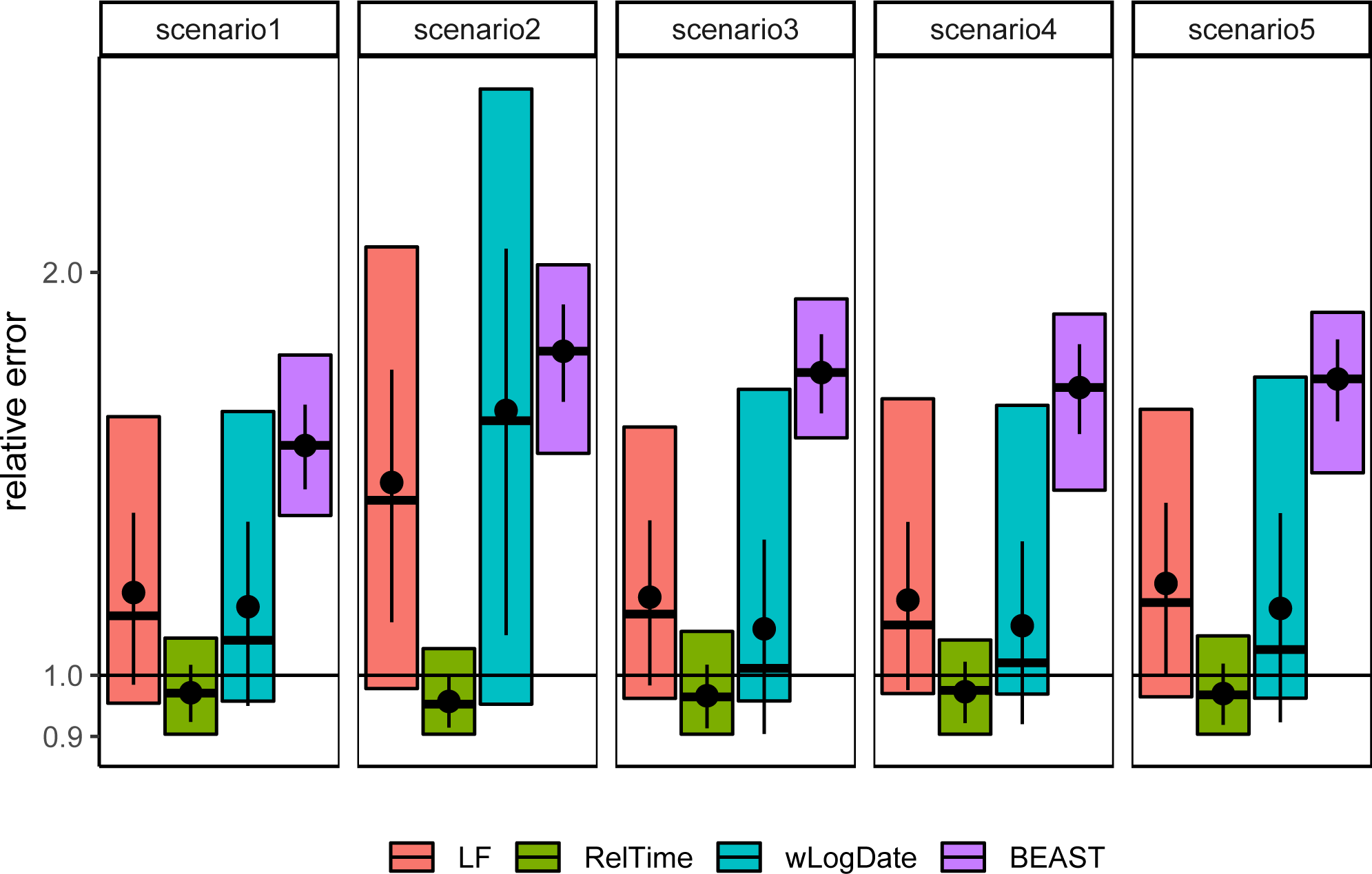
Relative error of wLogDate, RelTime, and LF on inferring the Angiosperm’s age on different settings of the simulation by [2]. Boxplots show median with 95% CI. Point ranges show mean with one standard deviation. Results for BEAST was obtained from the original study [2] on the same dataset but some different settings for the calibrations.

## References

[1] Akerborg, O., Sennblad, B., and Lagergren, J. 2008. Birth-death prior on phylogeny and speed dating. BMC Evolutionary Biology, 8(1): 77.

[2] Beaulieu, J. M., O’Meara, B. C., Crane, P., and Donoghue, M. J. 2015. Heterogeneous Rates of Molecular Evolution and Diversification Could Explain the Triassic Age Estimate for Angiosperms. Systematic Biology, 64(5): 869–878.

[3] Donoghue, P. C. J. and Yang, Z. 2016. The evolution of methods for establishing evolutionary timescales. Philosophical Transactions of the Royal Society B: Biological Sciences, 371(1699): 20160020.

[4] Drummond, A. J. and Rambaut, A. 2007. BEAST: Bayesian evolutionary analysis by sampling trees. BMC evolutionary biology, 7: 214.

[5] Drummond, A. J. and Suchard, M. A. 2010. Bayesian random local clocks, or one rate to rule them all. BMC Biology, 8(1): 114.

[6] Drummond, A. J., Ho, S. Y. W., Phillips, M. J., and Rambaut, A. 2006. Relaxed Phylogenetics and Dating with Confidence. PLoS Biology, 4(5): e88.

[7] Dudas, G., Carvalho, L. M., Bedford, T., Tatem, A. J., Baele, G., Faria, N. R., Park, D. J., Ladner, J. T., Arias, A., Asogun, D., Bielejec, F., Caddy, S. L., Cotten, M., D’Ambrozio, J., Dellicour, S., Di Caro, A., Diclaro, J. W., Duraffour, S., Elmore, M. J., Fakoli, L. S., Faye, O., Gilbert, M. L., Gevao, S. M., Gire, S., Gladden-Young, A., Gnirke, A., Goba, A., Grant, D. S., Haagmans, B. L., Hiscox, J. A., Jah, U., Kugelman, J. R., Liu, D., Lu, J., Malboeuf, C. M., Mate, S., Matthews, D. A., Matranga, C. B., Meredith, L. W., Qu, J., Quick, J., Pas, S. D., Phan, M. V. T., Pollakis, G., Reusken, C. B., Sanchez-Lockhart, M., Schaffner, S. F., Schieffelin, J. S., Sealfon, R. S., Simon-Loriere, E., Smits, S. L., Stoecker, K., Thorne, L., Tobin, E. A., Vandi, M. A., Watson, S. J., West, K., Whitmer, S., Wiley, M. R., Winnicki, S. M., Wohl, S., Wölfel, R., Yozwiak, N. L., Andersen, K. G., Blyden, S. O., Bolay, F., Carroll, M. W., Dahn, B., Diallo, B., Formenty, P., Fraser, C., Gao, G. F., Garry, R. F., Goodfellow, I., Günther, S., Happi, C. T., Holmes, E. C., Kargbo, B., Këita, S., Kellam, P., Koopmans, M. P. G., Kuhn, J. H., Loman, N. J., Magassouba, N., Naidoo, D., Nichol, S. T., Nyenswah, T., Palacios, G., Pybus, O. G., Sabeti, P. C., Sall, A., Ströher, U., Wurie, I., Suchard, M. A., Lemey, P., and Rambaut, A. 2017. Virus genomes reveal factors that spread and sustained the ebola epidemic. Nature, 544(7650): 309–315.

[8] Felsenstein, J. 1981. Evolutionary trees from DNA sequences: A maximum likelihood approach. Journal of Molecular Evolution, 17(6): 368–376.

[9] Fitch, W. M. 1971. Toward defining the course of evolution: Minimum change for a specific tree topology. Systematic Zoology, 20(4): 406–416.

[10] Forest, F. 2009. Calibrating the Tree of Life: fossils, molecules and evolutionary timescales. Annals of Botany, 104(5): 789–794.

[11] Gascuel, O. 1997. BIONJ: an improved version of the NJ algorithm based on a simple model of sequence data. Molecular Biology and Evolution, 14(7): 685–695.

[12] Gilbert, M. T. P., Rambaut, A., Wlasiuk, G., Spira, T. J., Pitchenik, A. E., and Worobey, M. 2007. The emergence of HIV/AIDS in the Americas and beyond. Proceedings of the National Academy of Sciences, 104(47): 18566–18570.

[13] Guindon, S., Dufayard, J.-F., Lefort, V., Anisimova, M., Hordijk, W., and Gascuel, O. 2010. New Algorithms and Methods to Estimate Maximum-Likelihood Phylogenies: Assessing the Performance of PhyML 3.0. Systematic Biology, 59(3): 307–321.

[14] Heath, T. A. 2012. A hierarchical Bayesian model for calibrating estimates of species divergence times. Systematic biology, 61(5): 793–809.

[15] Hedge, J., Lycett, S. J., and Rambaut, A. 2013. Real-time characterization of the molecular epidemiology of an influenza pandemic. Biology Letters, 9(5): 20130331.

[16] Hillis, D. M., Moritz, C., and Mable, B. K. 1996. Molecular Systematics, volume 2nd. Sinauer Associates.

[17] Ho, S. Y. 2014. The changing face of the molecular evolutionary clock. Trends in Ecology & Evolution, 29(9): 496 – 503.

[18] Ho, S. Y. W. and Phillips, M. J. 2009. Accounting for Calibration Uncertainty in Phylogenetic Estimation of Evolutionary Divergence Times. Systematic Biology, 58(3): 367–380.

[19] Ho, S. Y. W., Duchêne, S., and Duchêne, D. 2015. Simulating and detecting autocorrelation of molecular evolutionary rates among lineages. Molecular Ecology Resources, 15(4): 688–696.

[20] Huelsenbeck, J. P., Larget, B., and Swofford, D. 2000. A compound poisson process for relaxing the molecular clock. Genetics, 154(4): 1879–92.

[21] Jukes, T. H. and Cantor, C. R. 1969. Evolution of protein molecules. In Mammalian protein metabolism, vol. III (1969), pp. 21-132, volume III, pages 21–132.

[22] Kishino, H., Thorne, J. L., and Bruno, W. J. 2001. Performance of a Divergence Time Estimation Method under a Probabilistic Model of Rate Evolution. Molecular Biology and Evolution, 18(3): 352–361.

[23] Kodandaramaiah, U. 2011. Tectonic calibrations in molecular dating. Current Zoology, 57(1): 116–124.

[24] Kumar, S. and Hedges, S. B. 2016. Advances in Time Estimation Methods for Molecular Data. Molecular Biology and Evolution, 33(4): 863–869.

[25] Lalee, M., Nocedal, J., and Plantenga, T. 1998. On the implementation of an algorithm for large-scale equality constrained optimization. SIAM Journal on Optimization, 8(3): 682–706.

[26] Langley, C. H. and Fitch, W. M. 1974. An examination of the constancy of the rate of molecular evolution. Journal of Molecular Evolution, 3(3): 161–177.

[27] Lemey, P., Suchard, M., and Rambaut, A. 2009. Reconstructing the initial global spread of a human influenza pandemic: A bayesian spatial-temporal model for the global spread of h1n1pdm. PLoS currents, 1: RRN1031–RRN1031. 20029613[pmid].

[28] Lepage, T., Bryant, D., Philippe, H., and Lartillot, N. 2007. A General Comparison of Relaxed Molecular Clock Models. Molecular Biology and Evolution, 24(12): 2669–2680.

[29] Mai, U., Sayyari, E., and Mirarab, S. 2017. Minimum variance rooting of phylogenetic trees and implications for species tree reconstruction. PLOS ONE, 12(8): e0182238.

[30] Nee, S., May, R. M., and Harvey, P. H. 1994. The reconstructed evolutionary process. Philosophical Transactions of the Royal Society of London. Series B: Biological Sciences, 344(1309): 305–311.

[31] Nguyen, L. T., Schmidt, H. A., Von Haeseler, A., and Minh, B. Q. 2015. IQ-TREE: A fast and effective stochastic algorithm for estimating maximum-likelihood phylogenies. Molecular Biology and Evolution, 32(1).

[32] Pulqúerio, M. J. and Nichols, R. A. 2007. Dates from the molecular clock: how wrong can we be? Trends in Ecology & Evolution, 22(4): 180 – 184.

[33] Rambaut, A. and Grass, N. C. 1997. Seq-Gen: an application for the Monte Carlo simulation of DNA sequence evolution along phylogenetic trees. Bioinformatics, 13(3): 235–238.

[34] Rambaut, A. and Holmes, E. 2009. The early molecular epidemiology of the swine-origin a/h1n1 human influenza pandemic. PLoS currents, 1: RRN1003–RRN1003. 20025195[pmid].

[35] Rutschmann, F. 2006. Molecular dating of phylogenetic trees: A brief review of current methods that estimate divergence times. Diversity & Distributions, 12(1): 35–48.

[36] Rzhetsky, A. and Nei, M. 1993. Theoretical foundation of the minimum-evolution method of phylogenetic inference. Molecular Biology and Evolution, 10(5): 1073–1095.

[37] Saitou, N. and Nei, M. 1987. The neighbor-joining method: a new method for reconstructing phylogenetic trees. Molecular Biology and Evolution, 4(4): 406–425.

[38] Sand, A., Holt, M. K., Johansen, J., Fagerberg, R., Brodal, G. S., Pedersen, C. N. S., and Mailund, T. 2013. Algorithms for computing the triplet and quartet distances for binary and general trees. Biology, 2(4): 1189–209.

[39] Sanderson, M. J. 1997. A Nonparametric Approach to Estimating Divergence Times in the Absence of Rate Constancy. Molecular Biology and Evolution, 14(12): 1218–1231.

[40] Sanderson, M. J. 1998. Estimating rate and time in molecular phylogenies: beyond the molecular clock? In Molecular systematics of plants II, pages 242–264. Springer.

[41] Sanderson, M. J. 2002. Estimating Absolute Rates of Molecular Evolution and Divergence Times: A Penalized Likelihood Approach. Molecular Biology and Evolution, 19(1): 101–109.

[42] Sanderson, M. J. 2003. r8s: inferring absolute rates of molecular evolution and divergence times in the absence of a molecular clock. Bioinformatics, 19(2): 301–302.

[43] Schwartz, J. H. and Maresca, B. 2006. Do molecular clocks run at all? a critique of molecular systematics. Biological Theory, 1(4): 357–371.

[44] Shankarappa, R., Margolick, J. B., Gange, S. J., Rodrigo, A. G., Upchurch, D., Farzadegan, H., Gupta, P., Rinaldo, C. R., Learn, G. H., He, X., Huang, X. L., and Mullins, J. I. 1999. Consistent viral evolutionary changes associated with the progression of human immunodeficiency virus type 1 infection. Journal of virology, 73(12): 10489–502.

[45] Snir, S., Wolf, Y. I., and Koonin, E. V. 2012. Universal Pacemaker of Genome Evolution. PLoS Computational Biology, 8(11): e1002785.

[46] Tamura, K., Battistuzzi, F. U., Billing-Ross, P., Murillo, O., Filipski, A., and Kumar, S. 2012. Estimating divergence times in large molecular phylogenies. Proceedings of the National Academy of Sciences, 109(47): 19333–19338.

[47] Tamura, K., Tao, Q., and Kumar, S. 2018. Theoretical Foundation of the RelTime Method for Estimating Divergence Times from Variable Evolutionary Rates. Molecular Biology and Evolution, 35(7): 1770–1782.

[48] Tavaré, S. 1986. Some Probabilistic and Statistical Problems in the Analysis of DNA Sequences. Lectures on Mathematics in the Life Sciences, 17: 57–86.

[49] Thorne, J. L., Kishino, H., and Painter, I. S. 1998. Estimating the rate of evolution of the rate of molecular evolution. Molecular Biology and Evolution, 15(12): 1647–1657.

[50] To, T.-H., Jung, M., Lycett, S., and Gascuel, O. 2015. Fast Dating Using Least-Squares Criteria and Algorithms. Systematic Biology, 65(1): 82–97.

[51] Volz, E. M. and Frost, S. D. W. 2017. Scalable relaxed clock phylogenetic dating. Virus Evolution, 3(2). vex025.

[52] Volz, E. M., Koelle, K., and Bedford, T. 2013. Viral Phylodynamics. PLoS Computational Biology, 9(3).

[53] Wertheim, J. O., Fourment, M., and Kosakovsky Pond, S. L. 2012. Inconsistencies in Estimating the Age of HIV-1 Subtypes Due to Heterotachy. Molecular Biology and Evolution, 29(2): 451–456.

[54] Xia, X. 2018. DAMBE7: New and Improved Tools for Data Analysis in Molecular Biology and Evolution. Molecular Biology and Evolution, 35(6): 1550–1552.

[55] Xia, X. and Yang, Q. 2011. A distance-based least-square method for dating speciation events. Molecular Phylogenetics and Evolution, 59(2): 342 – 353.

[56] Zuckerkandl, E. 1962. Molecular disease, evolution, and genetic heterogeneity. Horizons in biochemistry, pages 189–225.

